# Engineering a Cell-Based Orthogonal Ubiquitin Transfer Cascade for Profiling the Substrates of RBR E3 Parkin

**DOI:** 10.1101/2024.09.14.613079

**Authors:** Shuai Fang, Li Zhou, Geng Chen, Xiaoyu Wang, In Ho Jeong, Savannah E Jacobs, Bradley R. Kossmann, Wei Wei, Jing Zhang, Geon H. Jeong, Ivaylo Ivanov, Angela M. Mabb, Hiroaki Kiyokawa, Bo Zhao, Jun Yin

## Abstract

The E3 ubiquitin (UB) ligase Parkin utilizes a Ring-Between-Ring (RBR) domain to mediate the transfer of UB to its substrates to regulate diverse cellular functions, including mitochondrial quality control, cell cycle progression, metabolism programming, and the establishment of synaptic functions. Mutations affecting the E3 ligase activity of Parkin are associated with cancer and Parkinson’s disease (PD). An essential role of Parkin is to synthesize UB chains on the surface of damaged mitochondria to initiate mitophagy. Still, it is not clear how Parkin carries out other biological functions through the ubiquitination of its downstream targets in the cell. We hypothesized that a comprehensive substrate profile of Parkin would facilitate the discovery of ubiquitination pathways underpinning its multifaceted roles in cell regulation and reveal mechanistic linkages between Parkin malfunction and disease development. Here, we used phage display to assemble an orthogonal ubiquitin transfer (OUT) cascade of Parkin that can exclusively deliver an engineered UB mutant (xUB) to Parkin and its substrates in living cells. We then generated a substrate profile of Parkin by purifying xUB-conjugated proteins from cells and identifying them by proteomics. The OUT screen identified Parkin substrates involved in DNA replication, protein translation, intracellular protein transport, and rhythmic regulation. Based on previous literature implicating alterations in membrane vesicle trafficking in PD, we verified Parkin-catalyzed ubiquitination of Rab GTPases (Rab1a, Rab5a, Rab5c, Rab7a, Rab8a, Rab10, an Rab13) as well as CDK5, with reconstituted ubiquitination reactions in vitro and in cells. We also found chemical-induced stimulation of mitophagy enhanced Parkin-mediated ubiquitination of Rab proteins. These findings demonstrate that the OUT cascade of Parkin can serve as an empowering tool for identifying Parkin substrates to elucidate its cellular functions.

## Introduction

Parkin is a Ring-Between-Ring (RBR) type of E3 ubiquitin (UB) ligase that regulates diverse cellular processes, including mitophagy, energy metabolism, cell cycle progression, and synaptic activity^1–6^. The best characterized role of Parkin is its engagement in a feedforward cycle with the PTEN Induced Kinase 1 (PINK1) to decorate damaged mitochondria with UB chains for mitophagy initiation^7–9^. In such a cycle, PINK1 is stabilized on the outer mitochondrial membrane (OMM) upon mitochondria depolarization and phosphorylates UB conjugated to the OMM proteins. Phosphorylated UB molecules then bind to Parkin and recruit it to the mitochondrial surface for PINK1-catalyzed phosphorylation that would release Parkin from an auto-inhibited state^10–13^. Activated Parkin synthesizes more UB chains on the mitochondrial surface for their phosphorylation by PINK1, which in turn recruits and activates more Parkin. Loss-of-function mutations in Parkin and PINK1 are the first and second most common cause of autosomal recessive Parkinson’s disease (PD), suggesting that defects in the mitochondria stress response may impinge on neurodegenerative processes associated with PD^14–15^. However, endogenous Parkin in neuronal cells was not able to induce mitophagy upon chemical damage of mitochondria, and the PINK1-Parkin pathway does not affect basal mitophagy in mice and Drosophila^16–19^. These findings suggest Parkin may regulate other cellular processes underpinning PD pathogenesis independent of mitophagy, and a profile of Parkin substrates would be instrumental in deciphering the role of Parkin in PD.

Parkin does not demonstrate strict substrate specificity in ubiquitinating OMM proteins for building UB chains on damaged mitochondria^2^. Still, the choice of early ubiquitination targets by Parkin on the OMM is important for mitophagy initiation. For example, Parkin ubiquitinates mitofusin 1 and 2 (MFN1/2), two transmembrane GTPases on the OMM, to signal their degradation by the proteasome^20–21^. The removal of MFN1/2 would stop mitochondria fusion so damaged mitochondria would not contaminate their healthy counterparts. Parkin also ubiquitinates Rho family GTPases Miro1 and Miro2 on the OMM and their induced degradation would freeze the movement of damaged mitochondria on the cytoskeleton to prepare for mitophagy^22–23^. Besides mitophagy, the PINK1-Parkin axis may also operate on cytosolic targets, such as PARIS, the transcriptional co-repressor, and affect mitochondrial biogenesis. Parkin-mediated ubiquitination and degradation of PARIS enhances the expression of PGC-1α to promote mitochondrial biogenesis, which has a protective effect on dopamine-producing neurons^24–26^.

Parkin can also be activated in the cytosol by binding to phosphorylated UB generated by PINK1 or through the oxidation of its Cys residues^27–29^. The non-mitochondrial function of Parkin plays key roles in tumor suppression by ubiquitinating proteins involved in cell proliferation, apoptosis, and metabolism^5^. For example, Parkin cooperates with CDC20 and CDH1, the subunits of a cell cycle-regulatory E3 known as the anaphase promoting complex (APC), to mediate the ubiquitination and degradation of mitotic regulators, such as Polo-like kinase-1, Aurora-A/B, and Cyclin B1, exerting anti-proliferative effects^30^. Parkin ubiquitinates and inhibits the pro-apoptotic protein BAK by preventing it from localizing to the mitochondria^31^, while Parkin stabilizes the anti-apoptotic protein Bcl-2 via monoubiquitination^31^. Both pathways suppress cancer cell survival ^32^. Parkin also ubiquitinates and inhibits the glycolysis enzyme pyruvate kinase M2 (PKM2), which suppresses the cancer cell-specific Warburg effect on metabolism^33^. These tumor suppressive actions of Parkin are consistent with the genetic and epigenetic inactivation of the Parkin gene found in a variety of human cancers^5, 34^. Furthermore, Parkin has been implicated in multiple diseases besides PD and cancer, including leprosy, cerebral ischemia, and autism spectrum disorder^6, 34–39^. Developing unbiased methods to profile Parkin substrates will facilitate the deconvolution of the pathogenic roles of Parkin.

Parkin, like other E3s, carries out the last step of UB transfer through the E1-E2-E3 enzymatic cascades to conjugate UB to the substrate proteins. There are more than 600 E3s encoded in the human genome^40–41^, including 14 RBR types of E3s^42^, and their combination with 2 E1s and ∼50 E2s^43–45^ constitute a complex network of UB transfer cascades and make it a significant challenge to identify the direct substrates of an E3 for the interpretation of its biological functions. To identify the substrates of a designated E3 in the cell, we constructed an orthogonal UB transfer (OUT) cascade of E3s consisting of engineered xE1-xE2-xE3 enzymes to exclusively transfer a UB mutant (xUB) through the xE3 to its ubiquitination targets for their identification by proteomics^46^. We have validated the OUT platform for profiling E3 substrates by constructing OUT cascades with HECT type E3s E6AP/UBE3A and Rsp5, and U-Box E3s CHIP and UBE4B, and identified their substrates important for cell metabolism, antiviral response, protein aggregation, and endoplasmic reticulum (ER)-associated stress^47–50^. In this study, we engineered the RBR domain of Parkin to extend the OUT cascade to RBR type E3s for profiling their cellular targets. Based on the substrate profile generated by OUT, we found potential roles of Parkin regulating membrane vesicle trafficking by verifying Parkin-catalyzed ubiquitination of a panel of Rab GTPases, including Rab1a, Rab5a, Rab5c, Rab7a, Rab8a, Rab10, and Rab13, that support vesicle trafficking in the cell. We also verified Parkin catalyzed ubiquitination of cyclin-dependent kinase 5 (CDK5), whose hyperactivation was implicated in neurodevelopment and neurodegenerative processes. We found Parkin promotes the degradation of these substrates in cells, and enhanced Parkin activity coupled with mitophagy induction stimulated the ubiquitination of the Rab proteins. Currently several studies have been reported in the literature on identifying Parkin substrates by following the change of protein ubiquitination levels with the enrichment of diGly-modified peptide or purification of ubiquitinated proteins by tandem UB-binding entities (TUBEs) ^9, 51–55^. The OUT cascade identified Parkin substrates by directly following xUB transfer from Parkin to its substrate proteins. It overcomes the cross-regulation between Parkin and other E3s in the cell that may indirectly affect the ubiquitination levels of the substrate proteins of other E3s. The construction and validation of the Parkin OUT cascade in this work provides an empowering tool for elucidating diverse Parkin functions in normal cell physiology and disease development.

## Results

### Design and construction of RBR libraries of Parkin for engineering the OUT cascade

The OUT cascade of Parkin was designed to exclusively transfer an engineered xUB with the R42E and R72E mutations to Parkin and then to its substrate proteins in the cell. The double mutations in xUB prevent its activation by the wildtype (wt) E1 (Uba1 and Uba6). Instead, xUB would be activated by xE1 (xUba1) with an engineered UB binding site and E2 binding site for engaging an engineered xE2 (xUbcH5 or xUbcH7) to transfer xUB from the xE1 to xE2^56^. xUbcH7 has the R5E and K9E mutations in its N-terminal helix that would match with the redesigned E2-binding site in the UB-fold domain (UFD) of xUba1^47^. We found xUB cannot be transferred from the xUba1-xUbcH7 pair to wt Parkin suggesting the mutated interfaces in the xUB∼xUbcH7 conjugate are incompatible with endogenous Parkin to mediate UB transfer (**Supplemental Figure S1**). The mutated residues in xUB and xUbcH7 are at different interfaces with Parkin. The residues in the N-terminal helix of UbcH7 would interact with a loop in the Ring1 domain of Parkin as shown by a modeled structure of human Parkin bound to the UbcH7∼UB complex based on NMR measurements and a crystal structure of fruit fly Parkin in a complex with UbcH7 ^57–58^ (**Figure 1A** and **Supplemental Figure S2A**). The equivalent sequence in the Ring1 domain of rat Parkin for interacting with the N-terminal helix of UbcH7 would be ^237^PCIACTD^243^ with residues P237, I239, A240, T242 and D243 constituting the binding interface, as shown by the crystal structure of rat Parkin^59^ (**Supplemental Figure S2, B-E**). We thus would thus randomize these residues in the RBR domain of rat Parkin for selecting mutants with restored activity with the xUB-xUbcH7 conjugate to mediate xUB transfer.

**Figure 1.**
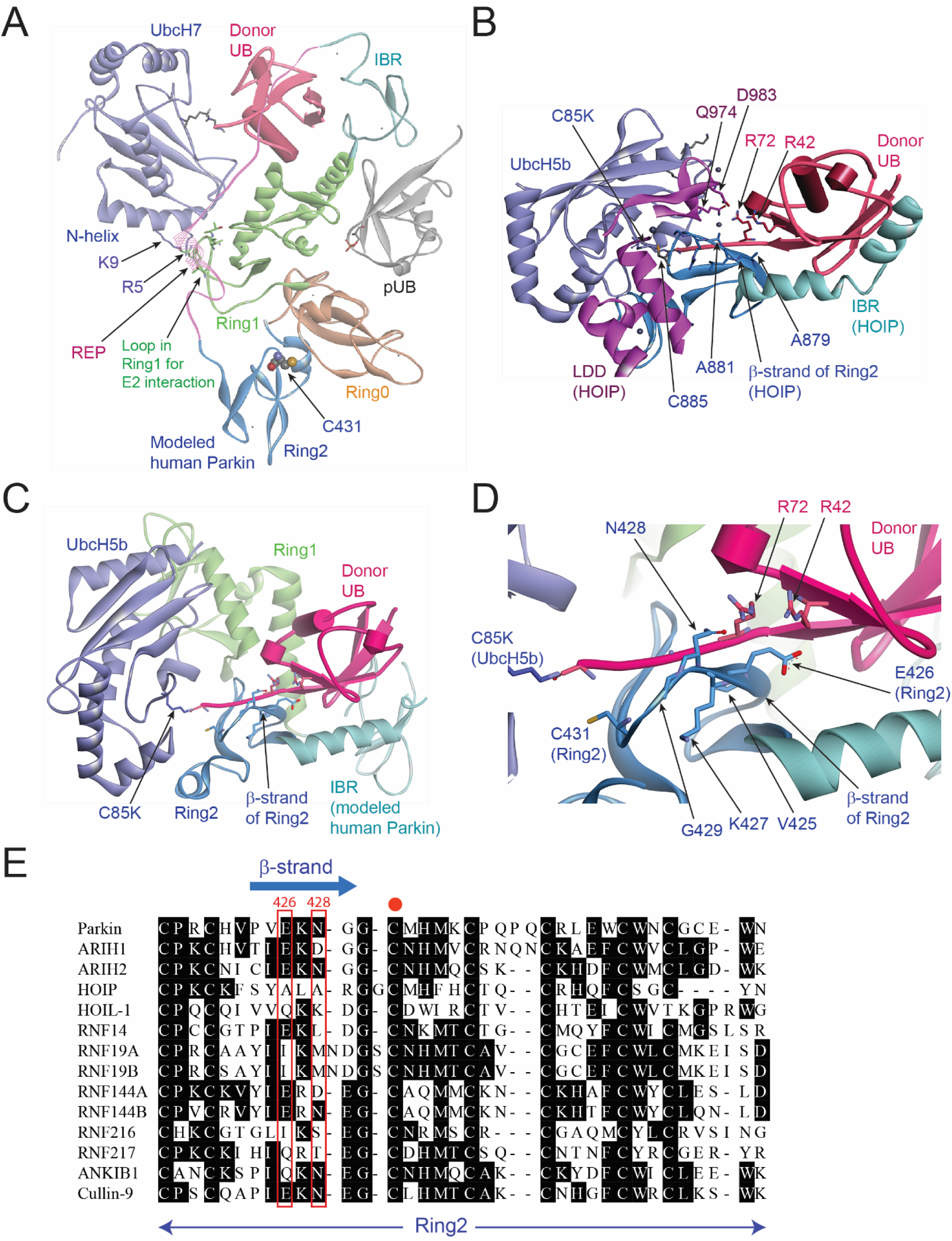
Structure analysis of the binding interface between Parkin RBR domain and the UB∼UbcH7 conjugate for engineering the OUT cascade with Parkin. (A) The modelled structure of human Parkin binding with the UbcH7∼UB conjugate (PDB ID: 6N13) ^58^. The modelled structure shows the binding between the N-terminal helix of UbcH7 and the loop region of the Ring1 domain of Parkin. (B) Crystal structure of HOIP RBR bound with the UbcH5b∼UB conjugate showing the UB C-terminus in an extended conformation and making contacts with the residues in the Ring2 and LDD domain of HOIP (PDB ID: 5EDV) ^60^. (C) Modelled structure of human Parkin RBR in a complex with the UbcH5b∼UB conjugate. The model was generated to mimic the structure complex between the RBR domain of HOIP and the UbcH5b∼UB conjugate in (B). (D) A detailed view of the interaction between the RBR domain of human Parkin with the donor UB bound to UbcH5b as shown in the modeled structure in (C). R42 and R72 of UB are in close vicinity to the residues constituting the β-strand with the sequence ^425^VEKNG^429^. (E) The alignment of the protein sequences of the Ring2 domain of the RBR E3s including Parkin and HOIP. Residues 426 and 428 (numbering in Parkin) in the β-strand of Ring2 for interacting with R42 and R72 of UB were highlighted in red frames. The catalytic Cys residue of the Ring2 domain is marked by a red dot.

The crystal structure of the RBR domain of HOIL-1-Interacting Protein (HOIP) with the UB∼UbcH5b conjugate revealed a binding interface between the R42 and R72 residues of UB and the Ring2 and LDD domain of HOIP (**Figure 1B**) ^60^. In the complex structure, the UB∼UbcH5b conjugate adopts an “open” conformation due to the tight engagement of Ring2 and LDD with the UB C-terminal residues ^71^LRLRGG^76^ that connect the catalytic Cys residue of UbcH5b with the globular domain of UB. The interactions between HOIP RBR and the UbcH5b∼UB conjugate also position the catalytic Cys (C885) in the Ring2 domain of the RBR within the attacking distance from the thioester bond of the UbcH7∼UB conjugate to enable the transthiolation reaction. The complex structure also showed that the R42 and R72 residues of UB, which are mutated to Glu in xUB, may engage LDD residues Q974 and D983 of HOIP through salt bridge and hydrogen bonding interactions. In addition, R42 and R72 of UB may interact with the β-sheet residues (^879^YALARG^883^) in the Ring2 domain of HOIP that runs in parallel to the C-terminal tail of UB (**Figure 1B**). Parkin does not have an LDD domain equivalent to HOIP, but the corresponding β-sheet residues in the Ring2 domain of Parkin has the sequence of ^425^IEKNG^429^ with residues E426 and N428 likely interacting with R42 and R72 of UB in the UB∼UbcH7 conjugate. To reveal the interaction between the Ring2 domain of Parkin and the UB∼E2 conjugate, we modeled the structure of the human Parkin RBR domain after the crystal structure of HOIP RBR with both RBRs bound with the UB∼UbcH5b conjugate (**Figure 1C**). In the modeled structure of the Parkin RBR, the β-strand containing the sequence ^425^VEKNG^429^ in the Ring2 domain of Parkin are indeed in contacting distance to R42 and R72 of UB and E426 and N428 of Parkin may engage the two Arg residues of UB with electrostatic and hydrogen bonding interactions (**Figure 1D**). We thus decided to construct a second rat Parkin library with randomized residues replacing I425, E426, N428, and G429 to restore the interaction between the RBR domain and xUB and enable xUB transfer through the engineered xParkin.

### Development of a phage display system for catalysis-based selection of the RBR domain of Parkin

We developed a phage display system for the selection of the RBR domain of Parkin based on its catalytic activity in pairing with xUbcH7 for the transfer of xUB (**Figure 2A**). In this system, the RBR domain library was expressed and anchored on the phage surface as an N-terminal fusion to the phage capsid protein pIII. Reaction of the RBR library with the xUba1-xUbcH7 pair would enable the transfer of biotin-xUB to the phage displayed RBR if the RBR variants had acquired complementary mutations to restore their interaction with the xUbcH7∼xUB conjugate. The formation of the RBR∼xUB conjugate would lead to biotinylation of the corresponding phage particles and they could be selected by binding to plates immobilized with streptavidin. Phage displaying catalytic active RBR domains were eluted by reaction with dithiothreitol (DTT) to cleave the thioester linkage between biotin-xUB and RBR. The eluted phage was then amplified by infecting *E. coli* cells for a subsequent round of selection. Through several rounds of phage selection, we expected to see the randomized residues in the RBR library to converge to a few consensus sequences to reveal the identity of catalytically active RBR variants that can assemble with the xUba1-xUbcH7 pair for xUB transfer.

**Figure 2.**
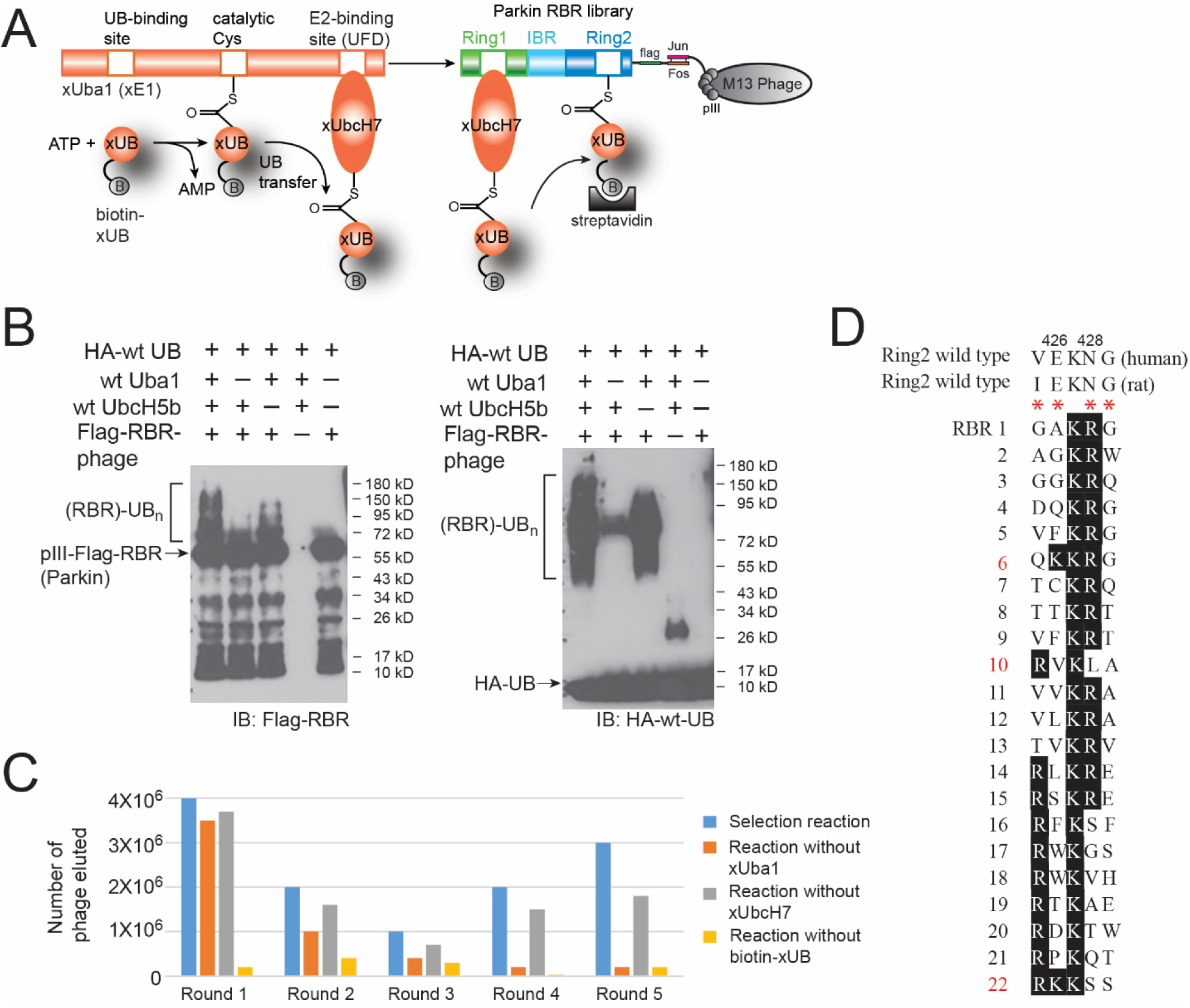
Phage selection of the RBR library of Parkin. (A) Selection of the RBR library of Parkin was based on the catalytic transfer of biotin-labelled xUB from xUba1 and xUbcH7 to the RBR variants displayed on the M13 phage. Once xUB is attached to the RBR domain on phage surface with the formation of the thioester conjugate, the corresponding phage particles are labelled by biotin and affinity selected by the interaction between biotin and immobilized streptavidin. (B) Reactivity of wt RBR domain of Parkin displayed on the phage particle. Ubiquitination reactions of the RBR domain on the surface of M13 phage were set up with HA tagged wt UB (HA-wt-UB), Uba1, UbcH7, and phage displaying the RBR domain. In control reactions, Uba1, UbcH7 or RBR displayed phage were excluded to establish the dependence of UB transfer to RBR through the cascade enzymes. (C) Selection of the RBR library of rat Parkin with randomized residues in the β-strand of the Ring2 domain. The phage library underwent five rounds of selection until clones from the selection showing convergent sequences at the randomized sites. In control reactions, xUba1, xUbcH7 or biotin-xUB was excluded from the reaction. Eluted phage titers from the selection reaction were greater than the control reactions, demonstrating the dependence of selection on the catalytic transfer of xUB from the xUba1-xUbcH7 pair to the RBR domain displayed on phage. (D) Alignment of the sequences of the RBR clones from the 5th round of selection of the Ring2 library, showing convergence of the selected clones in the randomized β-strand region. The randomized residues in Ring2 are designated by red stars.

To validate the phage selection strategy, we displayed the wt RBR domain of Parkin on the M13 phage and incubated the phage with wt Uba1, wt UbcH7 and biotin-wt UB to check the loading of UB on the RBR domain. In the presence of wt Uba1 and UbcH7, we observed strong ubiquitination of the RBR-pIII fusion displayed on phage and the UB loading on the RBR was dependent on Uba1 and UbcH7 (**Figure 2B**). In the absence of the E2 enzyme UbcH7, RBR could still be loaded with UB, but to a lesser extent, which was expected since E2-independent activity of RBR has been shown to support self-ubiquitination of RBR in reconstituted reactions^61^. After proving the wt Parkin RBR domain is functionally active when displayed on M13 phage, we carried out model selections for wt RBR displayed phage in parallel with a control phage that displayed an SV5V viral protein^62^. Both types of phage were reacted with wt Uba1 and UbcH7 to transfer biotin-wt-UB to the RBR. After the reaction, the phage was bound to the streptavidin plate and eluted with DTT. The phage titer showed that, with the same amount of phage input, the number of RBR phage retained by the streptavidin plate was more than 100-fold higher than that of the SV5V phage, demonstrating the high efficiency for selecting the RBR displayed phage based on the catalytic transfer of biotin-wt-UB through the Uba1-UbcH7 pair (**Supplemental Figure S3A**). We also carried out model selection for Parkin RBR displayed phage from mixtures with SV5V displayed phage. After the reaction of the phage mixtures with biotin-wt UB and the Uba1-UbcH7 pair, the phage mixtures were selected by binding to the streptavidin plate and the eluted phage particles were used to infect *E. coli* cells. We then used colony PCR to differentiate colonies infected with RBR or SV5V-displayed phage and found 8 out 10 clones selected from the 1/100 mixture of RBR/SV5V phage contained the pComb-RBR plasmid, and 1 out of 10 clones selected from the 1/1,000 mixture of RBR/SV5V phage contain the pComb-RBR plasmid (**Supplemental Figure S3B**). Therefore, each round of phage selection was able to enrich Parkin RBR phage by almost 100-fold, confirming the high efficiency of the phage display for engineering RBR variants based on UB transfer.

### Engineering the RBR domain of Parkin by phage display

We first carried out selection with the RBR library of Parkin with randomized loop residues in Ring1 that would interact with the N-terminal helix of UbcH7 (**Figure 1A and Supplemental Figure S2**). We reacted the phage library with biotin-xUB and the xUba1-xUbcH7 pair so biotin-xUB could be transferred to the RBR variants on phage if there was restored interactions. However, after five rounds of selection, we could not achieve convergence in the Ring1 library. These findings suggested that other interfaces such as that between the R42E and R72E mutations in xUB and the Ring2 domain of RBR could be important for restoring the interaction of RBR with the xUB∼xUbcH7 conjugate. We thus carried out phage selection with the RBR library randomized in the β-strand of Ring2 (**Figure 1C**). After five rounds of selection with increasing stringency, we observed the enrichment of phage particles from the selection reaction with the xUba1-xUbcH7 pair and biotin-xUB over controls (**Figure 2C**). In the fifth round of selection, we observed 20-30 fold higher number of phage particles eluted from the selection reaction compared to the controls, suggesting the phage selection was dependent on the catalytic transfer of biotin-xUB from the xUba1- xUbcH7 pair to the catalytic active RBR variants displayed on phage.

Sequencing of the Parkin RBR variants from the fifth round of selection revealed a strong preference for N428 to be mutated to a positively charged Arg or Lys residue within the sequence of ^425^IEKNG^429^ that was randomized in the Ring2 library (**Figure 2D**). In addition, the negatively charged residue Glu426 in the wt Parkin RBR was mutated to hydrophobic residues such as Val or Trp or positively charged residues like Lys or Arg. We picked several representative clones to screen their activity in mediating xUB transfer from the xUba1-xUbcH7 pair (**Figure 3A**). The selected RBR domains were expressed as GST fusions and the self-ubiquitination assay showed clone 6 with the sequence ^425^QKKRG^429^ and clone 22 with the sequence ^425^RKKSS^429^ had the highest activity in self-ubiquitination through xUB transfer from xUba1- xUbcH7. In contrast, clone 10 with the sequence ^425^RVKLA^429^ did not show strong activity of xUB transfer, suggesting positively charged Lys or Arg residues in position 426 or 428 in the Ring2 domain of the RBR were crucial for restoring its interaction with xUB. Based on the sequence alignment of the selected clones, there was not a significant consensus for the residues at position 425 which corresponded to an Ile in rat Parkin or a Val in human Parkin and at position 429 with a Gly in the native rat and human sequences (**Figure 2D**). We thus kept the native residues of I425 and G429 and incorporated the double mutations of E426R and N428R into the rat Parkin RBR domain. Generation of these mutations restored the self-ubiquitination of the engineered xRBR with xUB, suggesting the mutations could restore the binding of xUB in the xUbcH7∼xUB conjugate with the RBR domain (**Figure 3B**). In contrast, Parkin RBR with the single E426R mutation was less active with xUB transfer, suggesting both E426R and N428R mutations were needed to complement the R42E and R72E mutations in the xUB to enable the transfer of xUB to the xRBR domain. We next incorporated the E426R and E428R mutations into the RBR domain of human Parkin and found the mutant RBR can undergo efficient self-ubiquitination by uptaking xUB from the xUba1-xUbcH7 pair (**Figure 3C**). The full-length human Parkin with the same set of mutations in the RBR could also transfer xUB in the self-ubiquitination reaction (**Figure 3D**). Furthermore, human Parkin with the E426R and N428R mutations could transfer xUB to Miro1 (**Figure 3E**). Cumulatively, our results show that the Parkin mutant we engineered by phage display can function as an xParkin for the assembly of the OUT cascade to transfer xUB to Parkin substrates.

**Figure 3.**
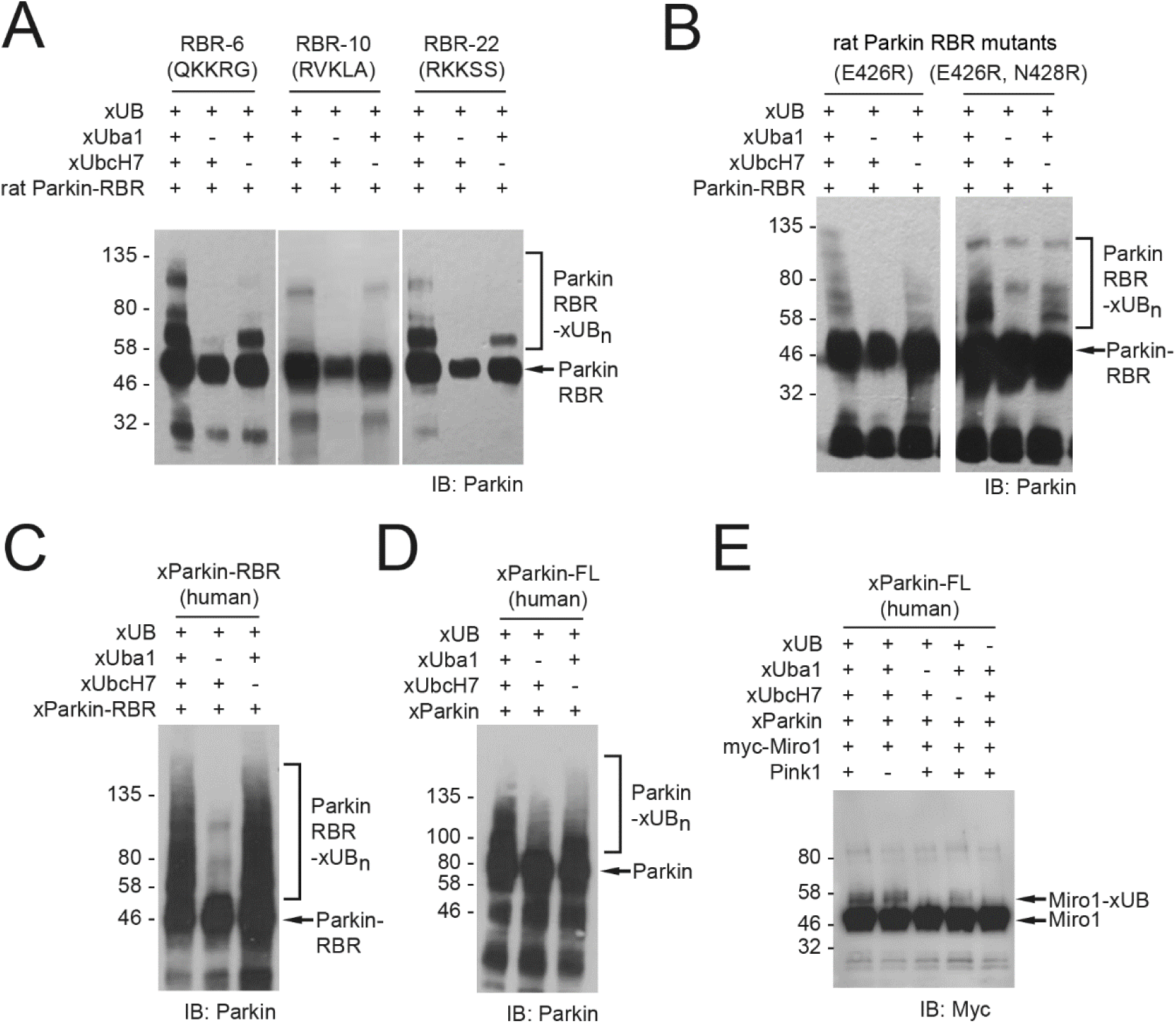
The xUB transfer activity of RBR variants from the selection of phage library with randomized Ring2 residues. (A) xUB transfer catalyzed by clones 6, 10 and 22 of the RBR mutants in the self-ubiquitination reaction. Clone 6 demonstrated the highest activity in xUB transfer. (B) Self ubiquitination activity of rat Parkin RBR with single mutation E426R and double mutation E426R/N428R. (C) The RBR domain of human Parkin with the E426R/N428R double mutations could mediate xUB transfer in the self-ubiquitination reaction. (D) The full-length (FL) human Parkin with the E426R/N428R double mutations also underwent self-ubiquitination with xUB. (E) Engineered human Parkin with the E426R/N428R double mutations could transfer xUB to known Parkin substrate Miro1.

### Profiling the Substrates of Parkin with the OUT cascade

The construction of the OUT cascade of Parkin gave us a platform for identifying Parkin substrates in cells. We previously screened HEK293 cells that would stably express the xUba1-xUbcH7 pair^47^. We co-transfected pLenti plasmids for the expression of HBT-xUB and myc-xParkin into HEK293 cells stably expressing the xUba1-xUbcH7 pair for expressing the OUT cascade of Parkin in the cell. To assay the expression of OUT components, we probed the OUT enzymes in the cell lysates by western blotting using antibodies against the specific tags fused to xUba1 (Flag), xUbcH7 (V5), and xParkin (myc) (**Figure 4A**). To confirm the activity of the OUT cascade in transferring xUB in the cell, we purified HBT-xUB conjugates from the cell lysate with the Ni-NTA resin and verified the presence of xUba1, xUbcH7 and xParkin in the resin-bound fraction. These results suggested the OUT cascade could be expressed in HEK293 cells and was active in transferring xUB to xParkin for its passage to Parkin substrates. We also generated the C431A mutant of xParkin with its catalytic Cys residue mutated to Ala and expressed the C431A mutant with the xUba1-xUbcH7 pair and HBT-xUB in HEK293 cells (**Figure 4A**). The C431 mutant was expressed at a similar level as the catalytic active xParkin in the cell, yet its presence in the fraction of HBT-xUB conjugated proteins bound to the Ni-NTA resin was decreased significantly, suggesting it only had very low activity in the transfer of xUB. Since the cells expressing the xUba1- xUbcH7 pair and the C431A mutant of xParkin would have a defective OUT cascade for xUB transfer, we used them as control cells to identify proteins that were preferentially labelled with HBT-xUB in cells expressing the catalytically active OUT cascade (OUT cells). In parallel runs, we purified xUB-conjugated proteins from the cell lysate of OUT cells and control cells using Ni-NTA and streptavidin affinity columns under denaturing conditions, and digested the proteins bound to the streptavidin resin with trypsin for proteomic analysis (**Figure 4B**).

**Figure 4.**
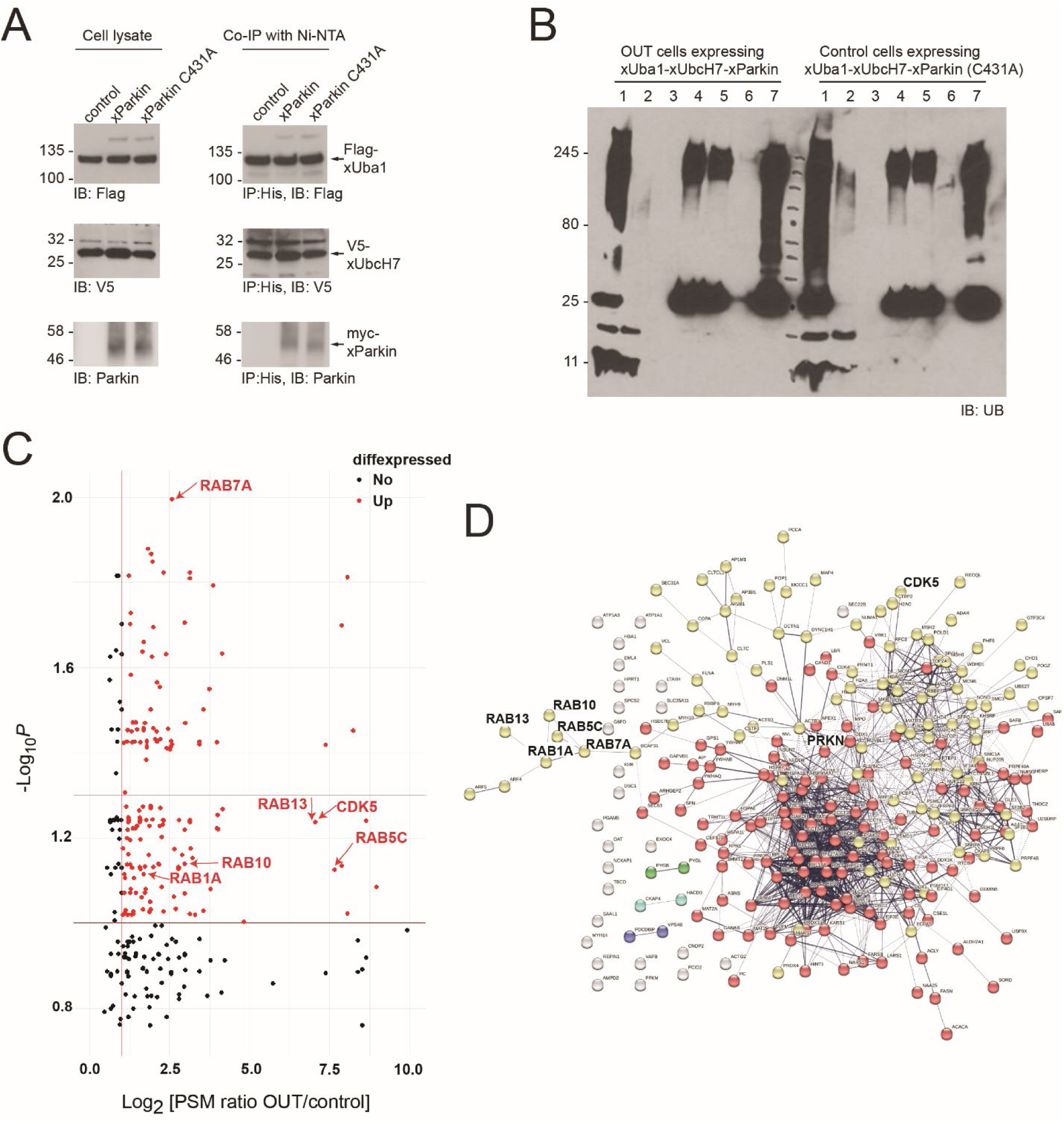
Expression of the OUT cascade of Parkin in HEK293 cells and identification of xUB-conjugated proteins. (A) Expression of the Parkin OUT cascade in HEK293 cells and confirmation of the activity of the OUT cascade in the transfer of xUB by assaying the presence of the OUT enzymes in the Ni-NTA pulldown samples. (B) Tandem purification of xUB-conjugated proteins from lysates of the cells expressing the catalytically active OUT cascade (OUT cells) and the control cascade with the C431A mutant of xParkin (control cells). Lane assignments: 1. cell lysates before Ni-NTA binding; 2. flow-through of the cell lysates after Ni-NTA binding; 3. wash of the Ni-NTA beads; 4. elution from the Ni-NTA beads; 5. flow-through from the streptavidin beads; 6. wash of the streptavidin beads; 7. proteins bound to the streptavidin beads. The western blot was probed with an anti-UB antibody to reveal ubiquitinated species enriched by tandem purification. (C) Volcano plot of Parkin substrates identified by the OUT screen. N = 3 independent biological replicates. Red dots designate proteins with Log_2_[PSM ratio OUT/control] >1 and -Log_10_*P* > 1. (D) Protein-protein interaction network for the targets identified from the OUT screen of Parkin substrates. The plot was generated from STRING (STRING PPI enrichment p-value < 1.0e-16). Line thickness indicates strength of data support.

We repeated the tandem purification and proteomic identification of xUB-conjugated proteins three times from Parkin OUT and control cells cultured in parallel to acquire the proteomic datasets for the identification of Parkin substrates (**Supplemental Table S1**). The volcano plot of the dataset identified 254 potential Parkin substrates with a *P* value < 0.1 (-Log_10_*P* > 1.0) and an average PSM ratio > 2 (Log_2_ [PSM ratio OUT/control] > 1) between the purification from the OUT and control cells. Among them, 104 proteins had a *P* value of medium stringency (*P* < 0.05, -Log_10_*P* > 1.3) and 27 proteins had a *P* value of high stringency (*P* < 0.01, -Log_10_*P* > 2) (**Figure 4C-4D, Supplemental Figure S4** and **S5**, and **Supplemental Table S2**). There were substantial overlaps (148 and 82 proteins) between the Parkin substrate profiles generated by the OUT screen and the Lys-ε-diGly peptide enrichment from previous studies ^55, 63^ (**Supplemental Table S3**). The OUT screen also identified many new Parkin substrates that have not been characterized before. Functional annotation-based bioinformatics analysis using DAVID showed that many substrates were involved in DNA replication, protein translation, rhythmic regulation, and intracellular protein transport **(Figure 5A-5C).** We also found numerous substrates that were assigned to the central nervous system development and the membrane vesicle-mediated transport clusters **(Figure 5D** and **Supplemental Figure S6A).** Moreover, disease-associated annotations related to Parkinsonism, neuropathy, neurodegeneration, intellectual disability, and glycogen storage disease were also identified (**Supplemental Figure S6B)**. Previous literature has demonstrated that disruptions in protein transport and vesicle trafficking mediated by Rab GTPases may play crucial roles in PD pathology^14, 64–66^. Therefore, we decided to verify Parkin catalyzed ubiquitination of the Rab proteins from the OUT screen. Besides Rabs, cyclin-dependent kinase 5 (CDK5) was also found to be a potential substrate of Parkin from the OUT screen. Hyperactivity of CDK5 has been associated with neurotoxicity observed in PD and it can suppress the E3 ligase activity of Parkin by phosphorylation^67–68^. We thus also proceeded to verify CDK5 as a Parkin substrate in follow-up ubiquitination reactions.

**Figure 5.**
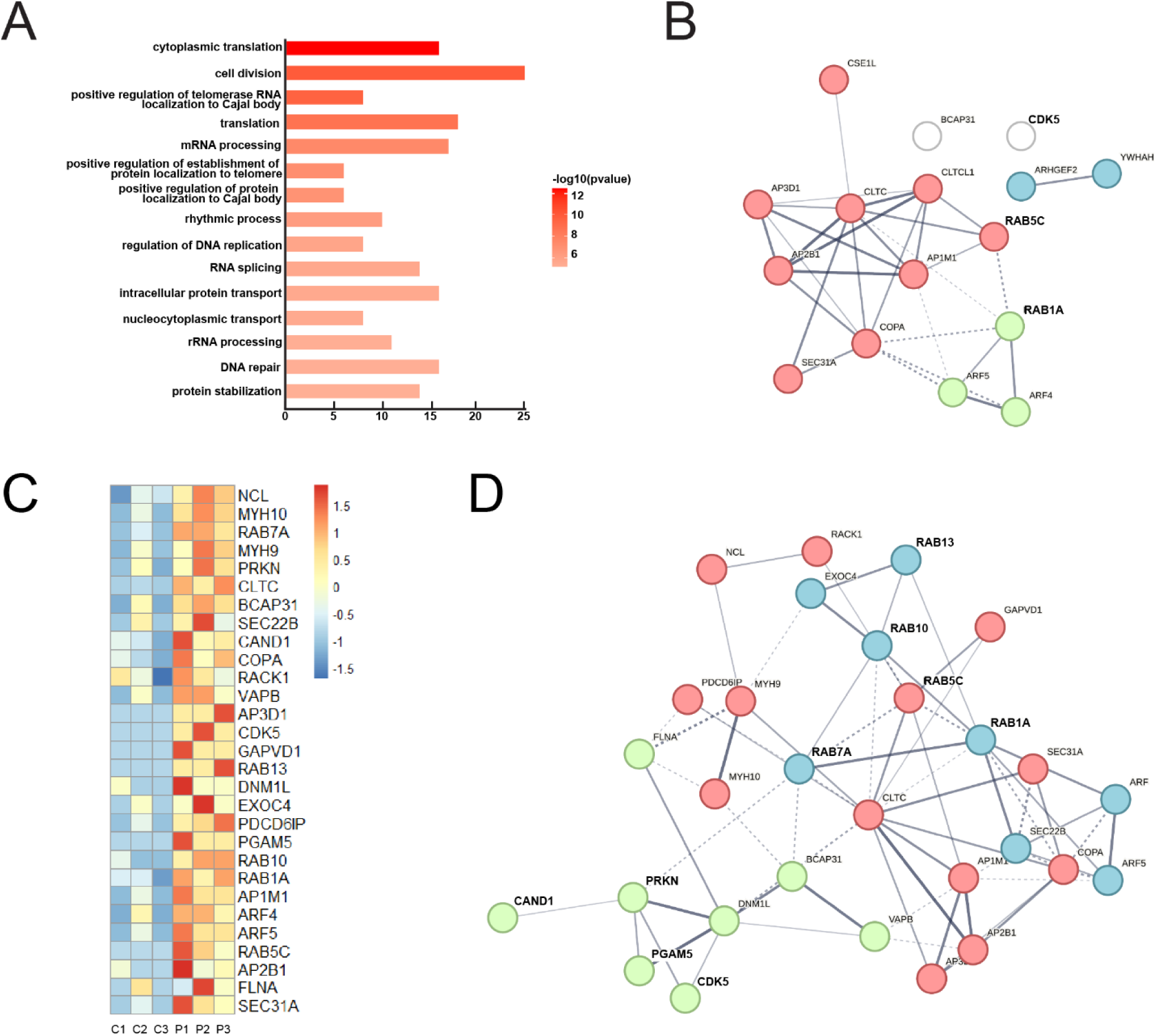
Analysis of the biological functions and interactions of Parkin substrates identified by the OUT screen. (A) Visualization of top significant terms from the list of gene ontology related with Biological Process. (B) STRING protein-protein interaction network for the targets identified from OUT that were classified under intracellular protein transport (GO:006886). (C) Heatmap with normalized PSM values for the targets identified from OUT that were classified under vesicle-mediated transport (GO:0016192). (D) STRING protein-protein interaction network for targets identified from OUT that were classified under vesicle-mediated transport.

### Validation of Parkin substrates by in vitro ubiquitination

The OUT screen identified Rab7a as a high-fidelity substrate with a -Log_10_*P* of 2.01 and Rab1a, Rab5c, Rab10, Rab13, and CDK5 with -Log_10_*P* in the range of 1.1 – 1.3. Rab8a was also identified in the OUT screen but with a nonsignificant *P* value. Rab5a was not identified by the OUT screen, but it is homologous to Rab5c from the OUT screen ^69^. We thus included Rab5a and Rab8a in the panel of Rabs for verification as Parkin substrates. We expressed and purified Rab GTPases from *E. coli* and set up in vitro ubiquitination reactions to verify their ubiquitination by wt Parkin (**Figure 6**). In the reconstituted reaction, we incubated the Rab proteins with wt UB and the cascade enzymes of Uba1-UbcH7-Parkin, and probed the formation of UB-substrate conjugates with a substrate-specific antibody. We also set up the in vitro ubiquitination reaction with the known Parkin substrate Mfn1 as a positive control^21^. We found Mfn1, along with Rab1, Rab5a, Rab7a, Rab10, and Rab13 could all be ubiquitinated by Parkin alone with the formation of substrate- UB conjugates, while the addition of PINK1 enhanced the ubiquitination of these substrates. Rab5c and Rab8a mainly showed mono-ubiquitination bands catalyzed by Parkin alone and the addition of Pink1 to the reaction mixture promoted the formation of polyubiquitinated Rab5c and Rab8a. These results verified Rab proteins as Parkin substrates, and as expected, PINK1 activation of Parkin enhanced the ubiquitination of Parkin substrates identified by the OUT screen. In addition, we set up an in vitro ubiquitination reaction of CDK5 with Parkin and found Parkin could ubiquitinate CDK5 alone, whereas similar to Rabs, the addition of PINK1 led to its enhanced ubiquitination.

**Figure 6.**
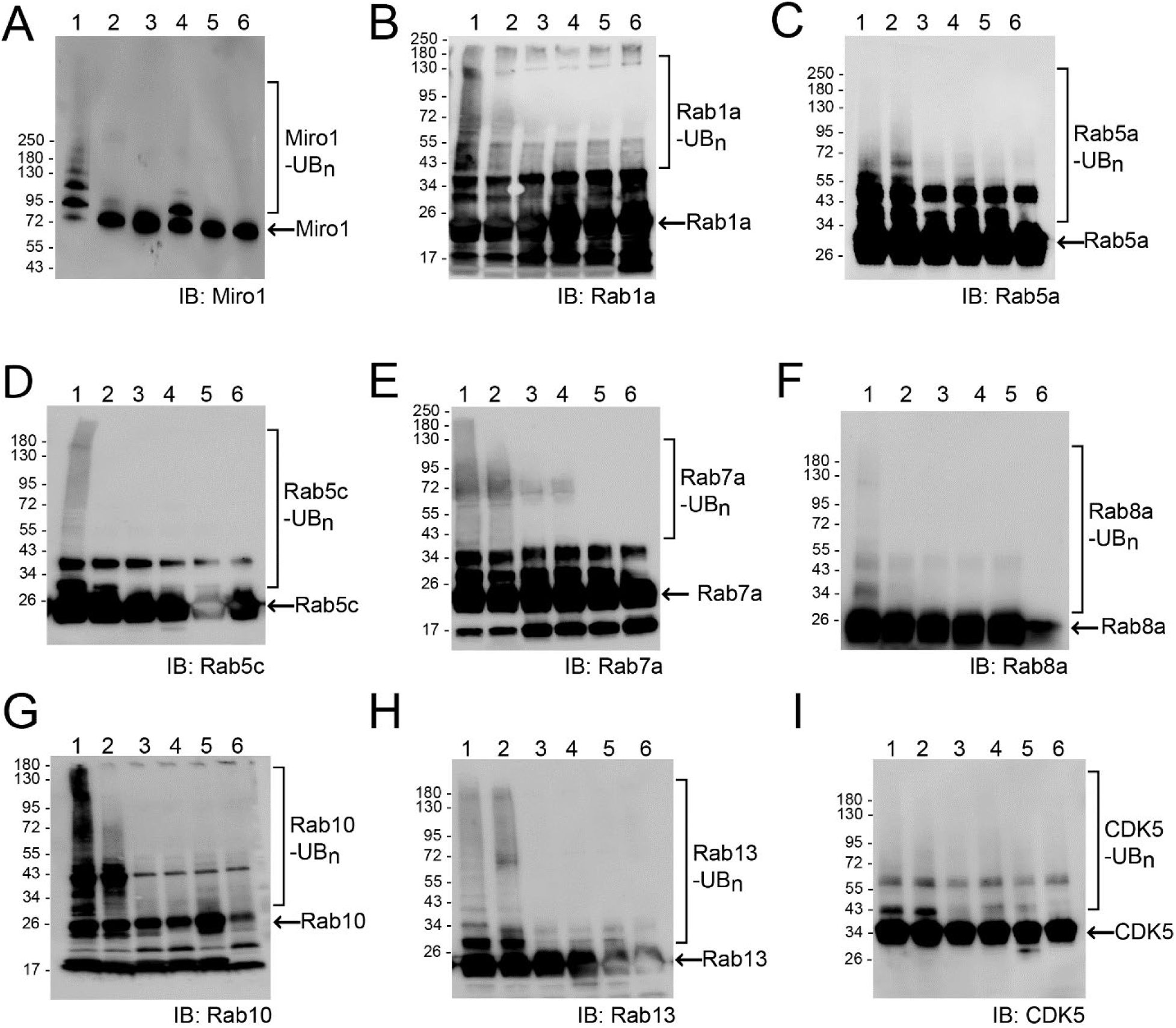
In vitro ubiquitination of Parkin substrates in reconstituted reactions. (A-I) Reactions were set up with ATP and wt Uba1, UbcH7 and Parkin for the transfer of wt UB to the Parkin substrates. Pink1 was also added to the reaction for the activation of Parkin by phosphorylating UB and Parkin. In control reactions, either Pink1 or each component of the UB transfer cascade of Parkin was excluded from the reaction to compare the activity of substrate ubiquitination in the presence of Parkin and Pink1. Lane assignments: 1, reaction with substrate proteins, PINK1 and the Uba1-UbcH7-Parkin cascade for the transfer of wt UB; 2, reaction missing PINK1; 3, reaction missing Uba1 as the E1; 4, reaction missing UbcH7 as the E2; 5, reaction missing Parkin as the E3; 6, reaction missing wt UB.

### Verification of Parkin-catalyzed substrate ubiquitination in HEK293 cells and the regulation of substrate stability by Parkin

We next verified the ubiquitination of identified Parkin substrates in the HEK293 cells. We transfected cells with an increasing amount of Parkin to enhance Parkin expression in the cell and treated the cells with MG132 to inhibit proteasome activity. We then immune purified substrate proteins and analyzed their ubiquitination. Increased Parkin expression in the cell enhanced substrate ubiquitination (**Figure 7**), demonstrating that Rab proteins and CDK5 are Parkin substrates. We also assayed the effect of Parkin-catalyzed ubiquitination on steady state levels of substrates in the cell by increasing Parkin levels. As Parkin expression increased, Rab GTPase and CDK5 levels decreased (**Figure 8**), suggesting Parkin regulates the steady state level of these substrates by ubiquitin-mediated degradation in the cell. If Parkin regulates the stability of the Rab proteins by ubiquitination, the suppression of Parkin activity in cancer cells may result in enhanced Rab expression. To verify this possibility, we examined the proteomic and genomic datasets available at the Clincal Proteomic Tumor Analysis Consortium (CPTAC) through the LinkedOmicsKB proteogenomics database (https://kb.linkedomics.org/). Our statistical analysis of the expression levels of Parkin, Rab1a, and Rab7a proteins in breast cancer tissues (n=88) indicated that there were significant reciprocal correlations between Parkin and Rab1a, as well as between Parkin and Rab7a. By contrast, no significant correlations were identified in the mRNA levels of Parkin with those of Rab1a or Rab7a in the same breast cancer tissues (**Supplemental Figure S7**). Therefore, Parkin expression is correlated inversely with Rab1a or Rab7a expression at the protein levels, not at the RNA levels, which is consistent with the posttranslational regulation of these Rab proteins by Parkin.

**Figure 7.**
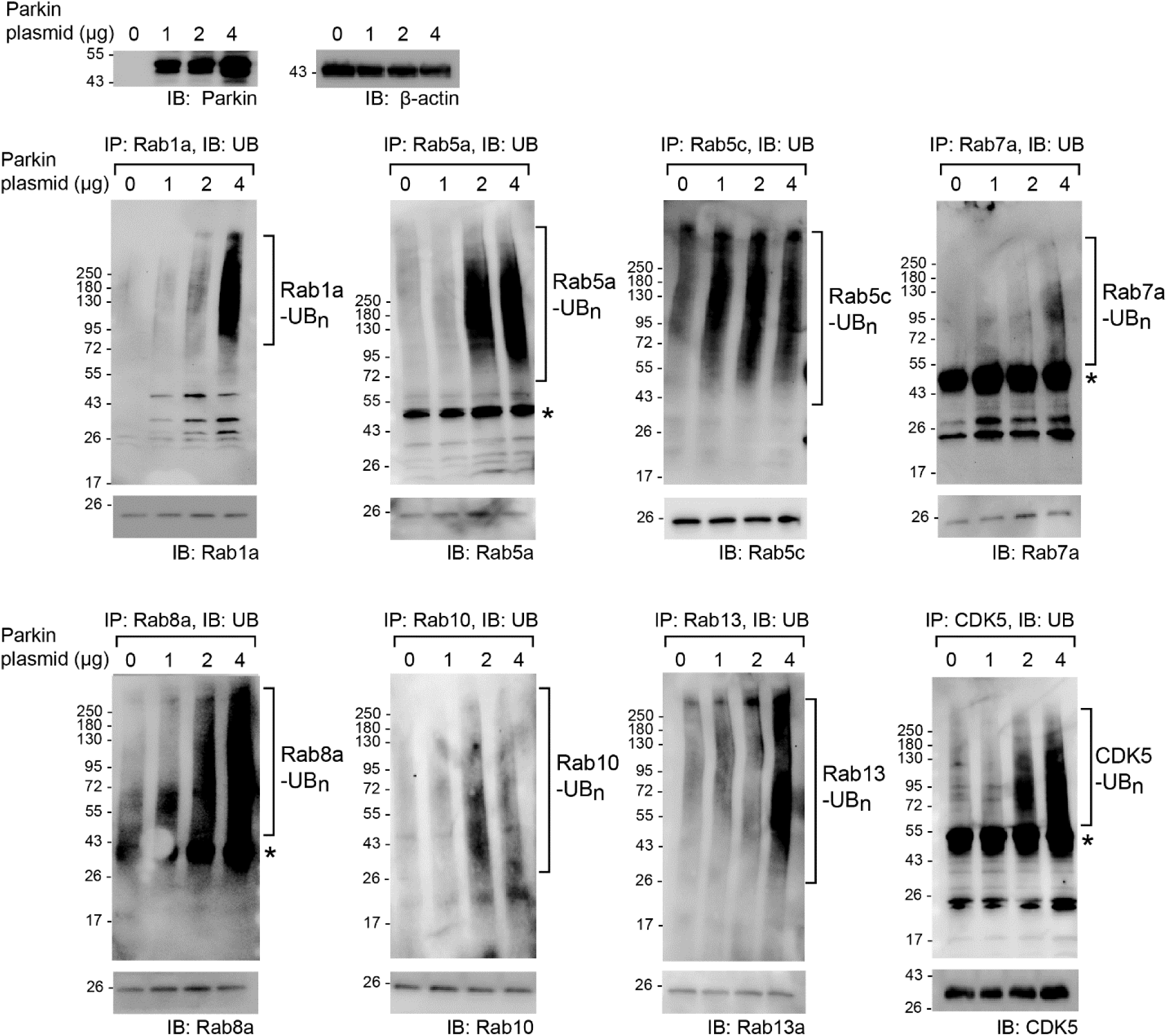
Validation of Parkin-induced ubiquitination of substrate proteins identified by the OUT cascade in cells. Parkin was over expressed in HEK293 cells with an increasing amount of Parkin plasmid used for cell transfection. The designated substrates were immune purified from the cell with specific antibodies and their ubiquitination levels were probed with an anti-UB antibody. Bands designated with a star on the western blots correspond to the size of IgG heavy chain.

**Figure 8.**
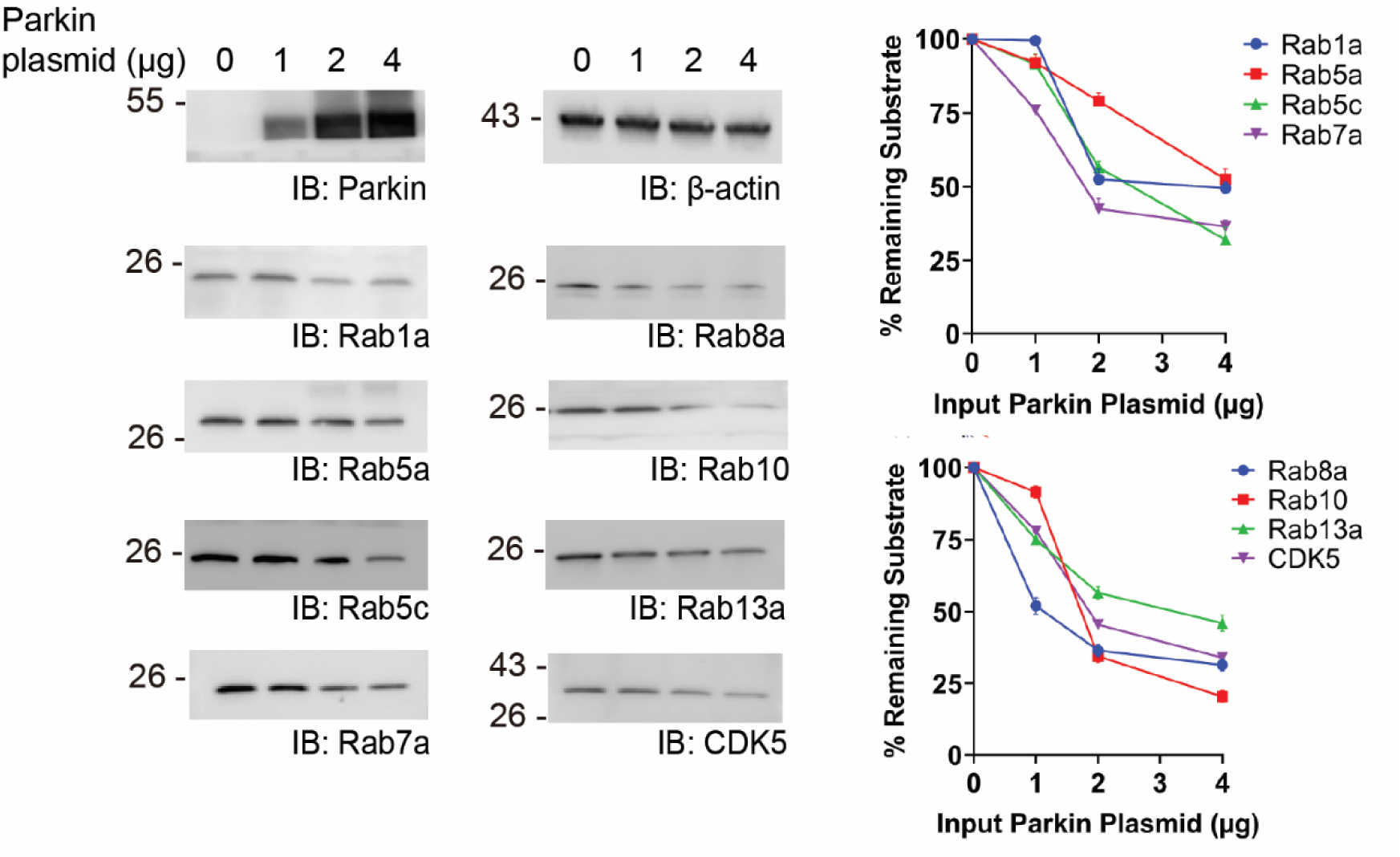
Accelerated degradation of Parkin substrates with increasing amount of Parkin expression. HEK293 cells were transfected with an increasing amount of Parkin plasmid and the level of designated substrates in the cell were assayed by western blots probed with substrate-specific antibodies. In the plots on the right, the relative intensities of the parkin substrate bands were quantified by the ImageJ software and plotted with the increasing amount of Parkin plasmid used for transfection. The levels of the substrate proteins in the cell were mean ± S.E. of three independent experiments with the vertical bars showing SEM of the experiments (n = 3).

### Enhanced ubiquitination of Parkin substrates with stimulated mitophagy

The stimulation effect of PINK1 on Parkin-catalyzed substrate ubiquitination in reconstituted reactions (**Figure 6**) prompted us to assay if Parkin activation by PINK1 during mitophagy induction in the cell would lead to enhanced ubiquitination of substrates identified by the OUT cascade. We added antimycin and oligomycin (AO) to HEK293 cells to induce mitochondrial depolarization^7^ and assayed the ubiquitination of Parkin substrates along with Mfn1 that is known to be ubiquitinated by Parkin to initiate mitophagy ^21^. As validation of the assay, Mfn1 underwent enhanced ubiquitination with AO stimulation or with forced Parkin expression or under both conditions. (**Figure 9**). Similarly, AO treatment significantly enhanced the ubiquitination of Rab1a, Rab5a, Rab7a and Rab10 in cells without forced expression of Parkin. AO treatment combined with increasing amounts of Parkin expression further increased the ubiquitination of Rab proteins. These results suggest that Parkin increases ubiquitination of Rab proteins upon the induction of mitophagy in the cell.

**Figure 9.**
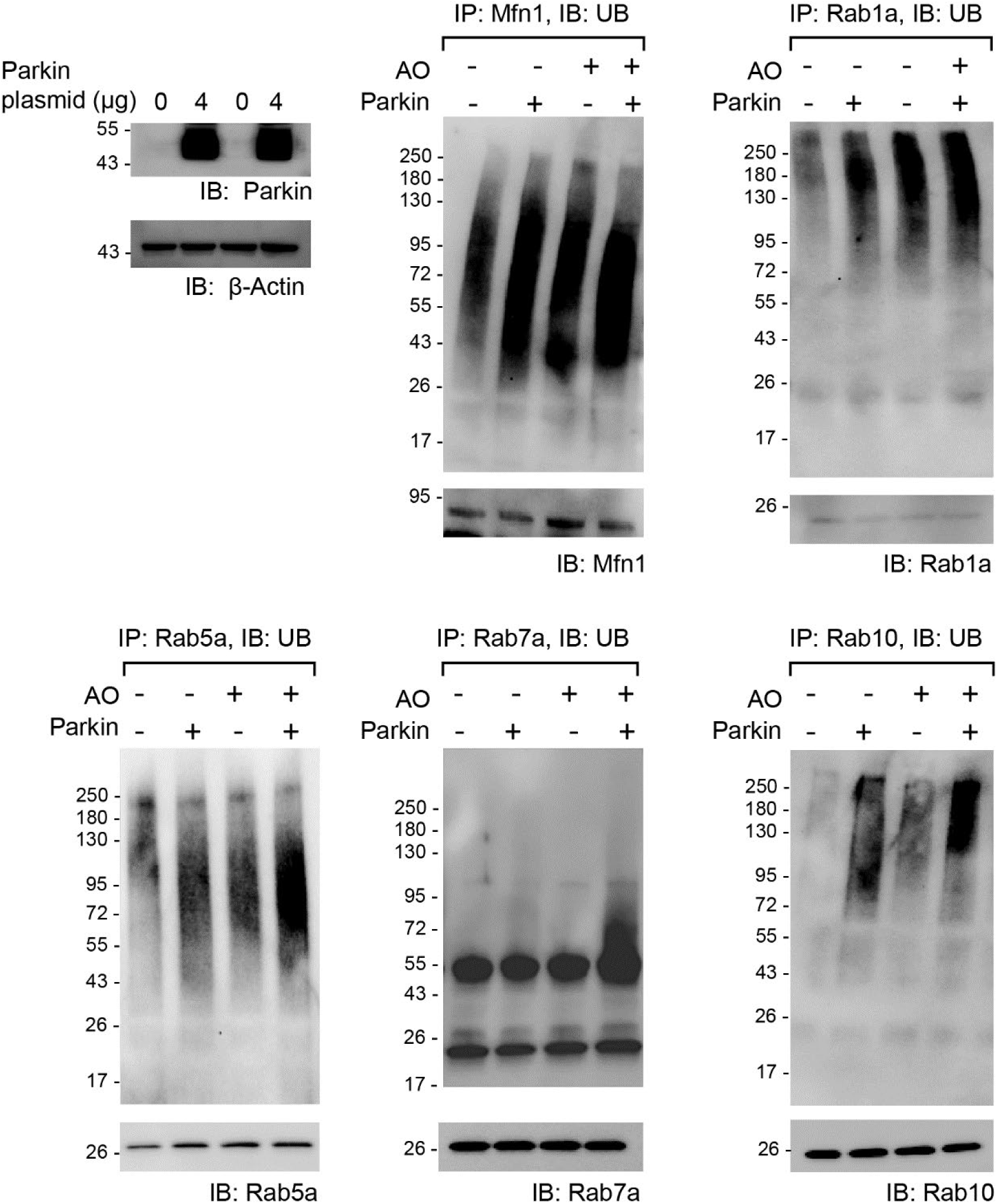
Induction of mitophagy led to enhanced ubiquitination of Parkin substrates. HEK293T cells with and without the transfection of the Parkin plasmid were treated with AO to induce mitophagy. Designated substrate proteins were immune purified with substrate specific antibodies and their ubiquitination levels were assayed by an anti-UB antibody.

## Discussion

In this study, we used phage display to engineer the RBR domain of Parkin so it can assemble an OUT cascade with the previously engineered xUba1-xUbcH7 pair for the exclusive delivery of HBT-xUB to Parkin and then to its substrates in the cell. Since the RBR domain of Parkin binds UbcH7 and UB in the form of a thioester conjugate, we identified two interfaces in the Parkin RBR for restoring the interaction with xUB∼xUbcH7. One interface is between the loop in Ring1 and the N-terminal helix of UbcH7 that harbors the R5E and K9E mutation in xUbcH7, and another interface is between the β-stand of Ring2 and the R42 and R72 of UB that were both mutated to Glu in xUB. Phage selection of the RBR library with randomized loop residues in Ring1 failed to enrich clones with converged sequences to mediate xUB transfer. This suggests the interactions between the N-terminal helix of UbcH7 and the Ring1 domain of RBR would not play a decisive role for the association of the Parkin RBR with the UB∼UbcH7 conjugate. Instead, as suggested by the crystal structure of HOIP RBR and the UB∼UbcH5b complex, the LDD and Ring2 domain of HOIP RBR would engage R42 and R72 of UB to force the UB to adopt an open conformation with respect to UbcH7 to enable the handover of UB to the RBR domain through thioester exchange. Our modeled structure of Parkin RBR with UB∼UbcH5b also confirmed key interactions between the β-stand of Ring2 and R42 and R72 of UB. Correspondingly, phage selection of the RBR library with randomized β-stand residues of Ring2 yielded consensus clones with restored activity for transferring xUB to Parkin RBR. Based on the phage selection results, we combined mutations E426R and N428R into the RBR domain of human Parkin to generate the engineered xParkin for the assembly of the OUT cascade. Thus, the combination of our modeling and phage selection results suggests R42 and R72 residues of UB constitute a key interface for the binding of UB∼UbcH7 conjugate with the catalytic active conformation of the RBR domain to mediate UB transfer to the RBR. Parkin has been shown to mediate E2-independent transfer of UB from E1 for self and substrate ubiquitination, suggesting the RBR domain of Parkin may directly bind to UB in the UB∼E1 conjugate to enable its transfer to the catalytic Cys residue of Parkin^61^. Furthermore, the Ring2 domain of Parkin itself can mediate the transthiolation reaction between the UB∼UbcH7 conjugate and the catalytic Cys on Ring2, suggesting the interaction between Ring2 and the UB∼UbcH7 conjugate is sufficient for the UB transfer reaction^70^. These studies match our observation that the interface between the Ring2 domain of Parkin RBR and the R42 and R72 residues of UB could be engineered for the transfer of xUB with the R42E and R72E mutations. Given the sequence of the β-strand region of the Ring2 domain of RBR E3s is highly conserved, it would be possible to transplant the E426R and N428R mutations from the Ring2 domain of xParkin to other RBR E3s for extending the OUT cascade to other RBR family members (**Figure 1E**).

The OUT screen identified 254 proteins as potential Parkin substrates (**Supplemental Table S2**). Among them, 23 were found to be mitochondrial proteins based on their overlap with the human MitoCarta database^71^ (**Supplemental Table S3**). Some of the mitochondrial proteins identified in the OUT screen are known Parkin substrates, including APEX1, HSD17B10 and VDAC2 ^55, 72–74^. So far, several studies have been reported on the identification of Parkin substrates in various cell types, including cancer cells and neuronal cells. The majority of these studies used an anti-Lys-ε-diGly antibody to enrich trypsinized peptides with diGly-Lys conjugate as a remnant of UB modification to reveal the difference in the protein ubiquitome with or without Parkin expression or mitophagy induction. Proteins with enhanced ubiquitination due to Parkin expression or activation during mitophagy were assigned as potential substrates of Parkin. In a pioneering study by Sarraf et al, proteins with enhanced ubiquitination were identified in HCT116, HeLa, and SH-SY5Y cells with Parkin expression and chemical stimulation of mitophagy to assemble a substrate profile of Parkin, in which 148 common substrates were found matching the Parkin substrate profile identified by OUT (**Supplemental Table S3**) ^55^. In another study by Agarwal et al, a Parkin substrate profile in prostate cancer PC3 cells were identified under the condition of enhanced Parkin expression without mitophagy induction, and it showed 82 common substrates with the Parkin substrate profile from the OUT screen ^63^ (**Supplemental Table S3**). The substantial overlap between the Parkin substrate profiles from the OUT screen and Lys-ε-diGly proteomics validates the use of the Parkin OUT cascade for substrate identification. A few other studies enriched Lys-ε-diGly peptides from neuronal cells with Parkin expression and mitophagy induction and a dozen or so substrates were identified to be overlapping with the Parkin substrate pool from our OUT screen ^53–54^. The smaller number of overlapping Parkin substrates identified in neuronal cells and HEK293 cells may suggest a greater difference in proteome composition between the two cell types. Another study identified Parkin substrates by Parkin-mediated transfer of wt UB in drosophila, and four overlapping substrates were identified when compared to the Parkin substrate profile from the OUT screen ^51–52^ (**Supplemental Table S3**).

Based on the OUT screen, we found Parkin can ubiquitinate a panel of Rab proteins, including Rab1a, Rab5a, Rab5c, Rab7a, Rab8a, Rab10 and Rab13, and regulate their stability in the cell. Rabs are GTPases that are prenylated at the C-terminus to anchor them on membrane structures, including plasma membrane, endosomes, ER, and Golgi^69^. Rab proteins define the identity of the cell membrane and program their functions in subcellular trafficking, sorting, recycling and degradation. Among the Rabs identified as Parkin substrates, Rab5a and Rab7a are endocytic Rabs that reside on early and late endosomes, respectively, to register them for degradation by fusion with the lysosome or for recycling by directing them to the trans-Golgi network (TGN) or the plasma membrane ^75–76^. Rab1a, Rab8a and Rab10 are exocytic Rabs that mediate anterograde transport of vesicles from the ER to the plasma membrane. Rab1a, Rab5a, Rab7a, and Rab10 have also been identified to play important roles in mitophagy. Upon PINK1 deposition to depolarized mitochondria to activate UB chain synthesis by Parkin, RABGEF1, a Rab5 GEF, binds to the UB chains conjugated to OMM proteins for recruiting Rab5 to damaged mitochondria. Then a Mon1-Ccz1 complex facilitates the conversion from Rab5 to Rab7 on the mitochondria surface and Rab7 engages the Atg9-bearing vesicles from the TGN or late endosome as the membrane source for autophagosome assembly^77–79^. Rab1 is anchored on the Atg9 vesicles for binding to the ULK complex and PI3KC3/VPS34 complex to prime the formation of the omegasome, the precursor of the autophagosome^80–82^. Furthermore, the recruitment of Rab10 to damaged mitochondria is dependent on PINK1 and Parkin activities. Rab10 promotes the association of the adaptor protein Optineurin (OPTN) with damaged mitochondria for mitophagy initiation^83^. Beyond autophagosome formation, Rab7a could bind to effector proteins such as RPIL for transporting autophagosomes to perinuclear regions for their fusion with the lysosome^84^. Also, Parkin-catalyzed ubiquitination on damaged mitochondria can tether the mitochondria with Rab5-anchored early endosomes that eventually fuse with the lysosome to undergo mitophagy-independent degradation^85^. In this study, we found Parkin to ubiquitinate a set of Rabs (Rab1a, Rab5a, Rab7a, and Rab10) that play crucial roles in mitophagy. We also found that the induction of mitophagy in the cell enhanced Parkin-catalyzed ubiquitination of the Rab proteins. These results suggest Parkin may affect Rab-mediated autophagosome assembly and lysosome processing besides its well-characterized role in UB chain synthesis on damaged mitochondria. Further work is warranted to reveal the effects of Parkin-catalyzed Rab ubiquitination on mitophagy.

Genetic analysis of PD patients substantiates the association of PD with deficits in endosome trafficking and mitophagy pathways in the cell and both pathways hinge on the correct functioning of a suite of Rabs^64–65^. For example, Rab5a and Rab7a are anchored on endosomes and endosomal maturation is accompanied by a Rab5a to Rab7a conversion, similar to the succession of the two Rabs on the autophagosome^86–88^. The subsequent trafficking and fusion of late endosomes with the lysosome are mediated by the Rab7a-RPIL complex. Both Rab5a and Rab7a were found to be regulated by ubiquitination during membrane trafficking. In fact, Rab7a has been found to be a Parkin substrate^89–90^ and Parkin catalyzed monoubiquitination of Rab7a at K38 can enhance the tethering between mitochondria and the lysosome in neuronal cells for the flux of amino acids to mitochondria for metabolic processing^91^. Monoubiquitination of Rab7a at K191 by an unidentified E3 was found to weaken the binding of RPIL with Rab7a and stall the fusion of the late endosome to the lysosome^92–93^. Similarly, monoubiquitination of Rab5 by an unknown E3 can affect the association of Rab5 with the endosome, decreasing receptor endocytosis^94^. Our results confirm Parkin can ubiquitinate both Rab5a and Rab7a, suggesting Parkin likely broadly alters endosomal trafficking in cells. Beyond mitophagy, it would be of interest to further characterize the role of Parkin in endosome regulation to reveal potential linkages between PD-associated Parkin mutations and endosomal malfunction.

The identification of Rabs as Parkin substrates may suggest Rabs as a nexus linking Parkin with other PD-causing genes such as *LRRK2* and *SNCA* encoding leucine-rich repeat kinase 2 (LRRK2) and α- synuclein, respectively. Rab5a, Rab8a and Rab10 are among the Rabs that are phosphorylated by LRRK2, whose mutation leads to enhanced activity in patients with familial PD^95–97^. Phosphorylation of membrane anchored Rab8a by LRRK2 enhanced the membrane deposition of LRRK2 that would increase its phosphorylation of Rab10 ^98^, and the phosphorylation of both Rabs by LRRK2 on the lysosome surface promotes secretion of lysosome contents for stress release ^99–100^. On the other hand, Rab8a and Rab10 with enhanced phosphorylation catalyzed by PD-related LRRK2 mutants showed increased binding with effector proteins (RILPL1 and RILPl2), which inhibit protein trafficking and hinder ciliogenesis and neuronal cell development^101–102^. Also, excessive phosphorylation of Rab8a and Rab10a due to LRRK2 mutation inhibits the binding of Rabs with the GDP dissociation inhibitor (GDI) that is responsible for extracting the GDP-bound Rab from the membrane. This would impair the proper recycling of Rab proteins in autophagy and lysosome pathways and may account for the accumulation of α-synuclein in PD^97, 103^. The PD-related LRRK2 mutation may also affect mitophagy through Rab10. It has been shown that LRRK2- mediated phosphorylation of Rab10 inhibits its interaction with the mitophagy adaptor OPTN, preventing OPTN recruitment to damaged mitochondria^83^. In addition, LRRK1, a paralog of LRRK2, and TBK1, a kinase phosphorylating OPTN to enhance its binding with the UB chains on mitochondria, have been found to phosphorylate Rab7a to facilitate the recruitment of Atg9-bearing vesicles for autophagosome assembly^104–106^. In our study, we found Parkin can ubiquitinate Rab5a, Rab7a, and Rab10 in response to mitophagy induction in the cell. Further studies on the cross-regulation of ubiquitination by Parkin and phosphorylation by LKKR1/2 and TBK1 on Rab proteins may reveal another layer of control for mitophagy initiation.

The membrane transport processes mediated by the Rab proteins also operate in neuronal cells to enable specialized functions, including neurite outgrowth, receptor signaling, and the transport and recycling of synaptic vesicles ^107–108^. Parkin is known to regulate the levels of dopamine transporter, AMPA, and NMDA receptors at the neuronal synapse as well as the trafficking and endocytosis of presynaptic vesicles^109–116^. So, Parkin-catalyzed Rab ubiquitination may underlie its regulation of synaptic activities. The Rab proteins verified as Parkin substrates in this study all have oncogenic potential, and their elevated expression levels have been found in various types of cancer cells. For example, Rab5 and Rab7 affects the endocytosis and intracellular transport of EGFR, and high levels of the two Rabs enable the cell to acquire invasive features such as focal adhesion disassembly and epithelial-mesenchymal transition ^117–121^. Rab13 controls protein trafficking pathways pertaining to cancer growth, including integrin recycling for cell migration and the membrane transport of GLUT4 and VGEFR for glucose uptake and angiogenesis ^122^. Our findings that Parkin mediates ubiquitination and degradation of these oncogenic Rabs, as well as the reciprocal correlations between Parkin and Rab1a or Rab7a in breast cancer tissues, reiterate the role of Parkin as a tumor suppressor. We also verified CDK5 as a Parkin substrate. The hyperactivation of CDK5 is linked to PD for its role in promoting oxidative stress and mitochondrial dysfunction in dopaminergic neurons, leading to neuronal inflammation and apoptosis ^123–125^. The identification of Parkin-mediated CDK ubiquitination and degradation matches with the neuroprotective role of Parkin that could be further verified.

Overall, our study developed an OUT cascade for profiling Parkin substrates in the cell. The verification of Parkin-catalyzed ubiquitination of the Rab GTPase panel and CDK5 from the OUT screen suggests potential roles of Parkin in regulating membrane vesicle trafficking and the associated pathways implicated in neurodegeneration. A limitation of our study is that we performed the OUT screen of Parkin substrates in HEK293 cells that are drastically different from neuronal or cancer cells in cellular function and proteome composition. Future work on applying the OUT cascade of Parkin to disease-relevant cell types may reveal pathogenic pathways directly related to Parkin-catalyzed substrate ubiquitination. Still, our study validated the OUT cascade of Parkin as an empowering tool to map E3 substrates and established a bridgehead for extending the OUT cascade to other RBR E3s to decipher their cellular functions.

## Methods

### Reagents and plasmids

pET-15b and pET-28a plasmids were from Novagen (Madison, WI, USA). pGEX-4T1 vector was from Amersham Biosciences (Piscataway, NJ, USA). pLenti4/V5-DEST-zeocin (K498000) and ViraPower Lentiviral Packaging Mix (K4975-00) were from Life Technologies (Carlsbad, CA, USA). The follow plasmids were from Addgene: pLenti-puro-ARID1A (39478), pEGFP-C1-Rab1a (49467), pcDNA5/FRT/TO-Flag-Rab5a (28043), pBS-L30-mRuby3-Rab5c(166426), pcDNA3-EGFP-Rab7a (28047), pEGFP-C2-Rab8a (86075), pEGFP-C1-Rab10 (49472), pEGFP-C1-Rab13a (49548), pGEX-2TK-CDK5 (24894), pRK-myc-human Parkin (17612). The plasmids for the expression of wt UB, Uba1, and UbcH7, and engineered xUB, xUba1, and xUbcH7 in *E coli* and mammalian cells were from a previous study^47^. The pComb3H plasmid for the display of the RBR domain was previously reported ^48^. XL1 Blue cells were from Agilent Technologies (Santa Clara, CA, USA). HEK293T cells were from American Tissue Culture Collection (ATCC) and cultured in high-glucose Dulbecco’s modified Eagles medium (DMEM) (Life Technologies, 11965092) with 10% (v/v) Fetal bovine serum (FBS) (Life Technologies, 11965092). Doxycycline, hygromycin, blasticidin, zeocin and puromycin were from GiBCO/Invitrogen (Carlsbad, CA, USA) and RPI (Mount Prospect, IL, USA). Anti-CDK5 antibody (sc-6247), Anti-myc antibody (sc-40), anti-Rab5a antibody (sc-166600), anti-Rab7 antibody (sc-376362), anti-V5 antibody (sc-271944), and anti-UB antibody (sc-8017) were from Santa Cruz Biotechnology (Dallas, TX, USA). Anti-Rab1a antibody (A14663), anti-Rab5c antibody (A20882), anti-Rab8a antibody (A20976), anti-Rab10 antibody (A4459), and anti-Rab13 antibody (A10571) were from ABclonal (Woburn, MA, USA). The antibodies were diluted between 200 and 500-fold to probe the western blots. Anti-Flag M2 antibody (F3165) was from Sigma-Aldrich (St. Louis, MO, USA) and diluted 1,000-fold for western blotting. The restriction enzymes for cloning were from New England Biolabs (Ipswich, MA, USA). The DNA oligonucleotides used for cloning in this study were listed in **Supplemental Table S4** and were ordered from Integrated DNA Technologies (San Diego, CA, USA).

### Construction of the protein expression plasmids

The genes of full-length human Parkin were amplified by polymerase chain reactions (PCR) with primers SF19 and SF20. The amplified fragments were digested by restriction enzymes BamHI and NotI and cloned into pGEX-4T1 vector. The genes of the Parkin RBR domain were amplified by PCR with primers SF19 and SF30. The amplified fragments were also digested with BamHI and NotI and cloned into the pGEX-4T-1 plasmid. For the expression of full-length Parkin with mutated RBR domains, mutant RBR genes were amplified with primers SF19 and SF20 and cloned into the pGEX-Parkin vector between restriction sites BamHI and NotI. To characterize the mutant Parkin RBR domains from phage selection, the genes of the mutant RBR domains were amplified by PCR from the pComb vectors with primers SF19 and SF30. The amplified DNA fragments were digested by BamHI and NotI, and cloned into pGEX vector. The gene of full-length Parkin was cloned into pLenti-vector between the BstbI and NheI restriction sites with primers SF19 and SF20. The Parkin mutant with E426R and N428R mutations were introduced into wt Parkin by primers SF19, SF22, SF23, SF20 with overlapping PCR. The assembled PCR fragments were digested by BstbI and NheI and cloned into the pLenti vector. To construct the inactive Parkin mutant, the C431A mutation was introduced into pLenti-xParkin by primers SF19, SF22, SF31, SF20 with overlapping PCR. For the expression of Parkin substrates, the CDK5 gene was amplified from pGEX-CDK5 plasmid with primers SJ1 and SJ2, Rab1a amplified with SJ3 and SJ4, Rab5a with SJ5 and SJ6, Rab5c with SJ7 and SJ8, Rab7a with SJ9 and SJ10, Rab8a with SJ11 and SJ12, Rab10a with SJ13 and SJ14, and Rab13 with SJ15 and SJ16. The PCR fragments were digested with SacII/NotI and cloned into the pET-28a plasmid. The pET and pGEX plasmids were transformed into BL21(DE3) pLysS chemical competent cells (Invitrogen) for protein expression.

### Constructing a model of the human Parkin RBR-UbcH5b-UB complex

For modeling the human Parkin-UbcH5b-UB complex, we employed the structure of HOIP-RBR/UbcH5b-UB (PDB ID: 5EDV)^60^. Initially, homology models of the human Parkin dimer (residues 228-378 and 393-464) were generated using the structure of HOIP RBR (PDB ID: 5EDV)^60^ as a template. The model was then superimposed onto the 5EDV structure to obtain the human Parkin-UbcH5b-UB complex. Finally, structural minimization, including the adjustment of side chains, was performed to optimize their orientations using the Rosetta Relax protocol implemented in ROSETTA^126^.

### Displaying the Parkin RBR domain on M13 phage

The RBR domain of the rat Parkin was amplified by PCR with primers (GC1 and GC2) and cloned into the phagemid vector pComb3H between the SacII and SpeI restriction sites to display the RBR on phage with an N-terminal Flag tag and as a fusion to the phage pIII protein at the C-terminus. The preparation of the RBR displayed phage was performed following the protocol reported previously ^48^. Briefly, *E. coli* SS320 cells were transformed with the pComb3H phagemid vector, cultured in the 2×YT media, and infected with M13KO7 helper phage. The infected cell culture was shaken at 37°C overnight in 100 mL 2×YT that was supplemented with 100 µg/mL ampicillin and 50 µg/mL kanamycin. The next day, the cells were pelleted by centrifugation, and the phage particles in the supernatant were PEG precipitated and resuspended in TBS buffer (20 mM TrisHCl, 150 mM NaCl, pH 7.5). Phage titrations were performed with *E. coli* XL1- blue cells with standard procedures ^127–128^. Phage displaying Parkin library for selection and SV5V for model selection were prepared following the same procedure. The display of the RBR domain on phage was confirmed by western blotting probed with an anti-Flag antibody. To confirm the catalytic activity of the RBR domain displayed on phage surface, ubiquitination reactions were set up with 2 × 10^10^ phage particles, 0.5 μM Uba1, 5 μM UbcH5b, and 20 μM HA-wt UB in a reaction buffer with 50 mM tris-HCl (pH 7.5), 10 mM MgCl_2_, 5 mM ATP, and 50 μM dithiothreitol (DTT). After 1-hour incubation at 37°C, the reaction mixture was analyzed by SDS–polyacrylamide gel electrophoresis (PAGE) and western blotting probed with an anti-HA antibody to detect the conjugation of HA-UB to the RBR domain on phage or with an anti-Flag antibody to detect the ubiquitinated RBR-pIII fusion.

### Model selection of phage displayed Parkin RBR

UB transfer reaction with phage displayed Parkin was set up in a total volume of 100 μL with 0.5 μM wtUba1, 5 μM wtUbcH7, and 5 μM biotin–wt UB in a reaction buffer with 50 mM tris-HCl (pH 7.5), 10 mM MgCl_2_, 5 mM ATP, and 50 μM dithiothreitol (DTT), and 1 × 10^11^ phage with Parkin and SV5V displayed phages mixed at a ratio of 1/10, 1/100, or 1/1000 in TBS buffer (pH 7.5). The reaction was allowed to proceed for 1 hour at room temperature before it was added to 400 μL 3 % BSA in TBS buffer (pH 7.5). 100 μL of phage solution was distributed to each well of a 96-well plate coated with streptavidin. The streptavidin plate was incubated at room temperature for 1 hour. The supernatant was discarded, and the plate was washed 30 times with 0.05% (v/v) Tween 20, 0.05% (v/v) Triton X-100 in TBS and 30 times with TBS, each time with 200 µL of solution per well. After washing, phage bound to the streptavidin surface were eluted by adding 100 μL of 10 mM dithiothreitol (DTT) in TBS. Eluted phage were combined, added to 10 mL of log phase E. coli XL1-Blue cells and shaken at 37°C for 1 hour to infect the cells. The cells were then plated on LB agar plates supplemented with 2% (w/v) glucose and 100 μg/mL ampicillin. After overnight incubation at 37°C, colonies on the plates were analyzed by colony PCR with primers Jun13 and Jun14. To set up colony PCR, 40 µL reaction mixture for each PCR reaction was prepared, containing 29.4 µL H_2_O, 4 µL Mg^2+^ free PCR buffer (Promega), 4 µL 25 mM MgCl_2_, 1 mL primer Jun13 (10 µM), 1 µL primer Jun14 (10 µM), 0.32 µL dNTP mix (25 mM each dNTP) and 0.2 µL Taq DNA polymerase (Promega). Pipette tips were used to transfer colonies from the agar plates to the PCR reaction mixture. The PCR reactions were run with the following program: 95°C for 10 minutes, then 35 cycles of 94°C for 1 minute, 50°C for 45 seconds, 72°C for 1.5 minute and a final step of 72°C for 5 minutes. For analysis, 10 µL of the PCR reaction was loaded on a 1% agarose gel and the PCR products were separated by electrophoresis.

### Construction of the RBR library of Parkin for phage display

Residues to be randomized in the Parkin RBR (I425, E426, N428, and G429) were first mutated to Ala to generate phagemid pComb-Parkin-RBR-4Ala. For mutagenesis, the RBR gene was amplified by two sets of primers GC1/GC3 and GC4/GC2. The PCR fragments were assembled by overlap extension for cloning into the pComb phagemid between restriction sites SacII and SpeI. For randomization of the designated residues in the Parkin RBR, pComb-Parkin-RBR-4Ala was used as the template for PCR reactions with primer pairs GC1/GC3 and GC5/GC2. The overlap extension of the amplified fragments was cloned into pComb phagemid between restriction sites SacII and SpeI to generate the phagemid library. The library DNA was transformed into XL1-Blue cells by electroporation. The cells were plated on LB-agar plates containing ampicillin (100 μg/ml) and glucose (2%) and incubated overnight at 37°C. A total of 10 transformation reactions were performed, each time with 50 µL of competent cells. The transformation efficiency of the library DNA was titered and the size of the library was estimated to be 2× 10^7^, exceeding the theoretical diversity of the Parkin library with four randomized sites (1.6 × 10^5^). The cells growing on the plate were scrapped and library phagemid DNA was prepared with a DNA Maxiprep kit (Qiagen).

### Phage selection of the Parkin RBR library

For preparing phage displaying the Parkin library, 1 µL of the library phagemid DNA was used to transform 100 µL electrocompetent XL1Blue cells. After electroporation, the cell culture was shaken at 37°C overnight in 100 mL 2×YT that was supplemented with 100 µg/mL ampicillin and 50 µg/mL kanamycin. The next day, the cells were pelleted by centrifugation, and the phage particles in the supernatant were PEG precipitated and resuspended in TBS buffer (20 mM TrisHCl, 150 mM NaCl, pH 7.5). Phage titrations were performed with E. coli XL1- blue cells with standard procedures ^127–128^. For the first round of phage selection, 10^10^ phage particles were reacted with 0.5 μM xUba1, 10 μM xUbcH7, and 0.5 μM biotin-xUB for 1 hour. Reactions were diluted 10-fold into 0.1% bovine serum albumin (BSA)– TBST (20 mM TrisHCl, 150 mM NaCl, 0.1% Tween 20, pH 7.5) and bound to a streptavidin plate for 1 hour at room temperature. The plate was washed 30 times with 0.1% BSA-TBST, and then 100 µL TBS containing 100 mM DTT was added to each well to elute the phage particles. Each 100 µL of the phage elution solution was added to a culture of 1 mL XL1-Blue cells at the log phase of growth, and the culture was shaken slowly for 2 hours at 37°C to enable phage infection of the cell. Then the cells were plated onto LB-agar plates containing carbenicillin (100 μg/ml) and glucose (2%). After overnight incubation at 37°C, the colonies on the plates were collected, and the phagemid DNA was extracted with a DNA Miniprep kit. In parallel, control reaction was set up with xUba1, xUbcH7 or biotin-xUB missing from the reaction mixture. Phage in the control reactions were also bound to the streptavidin plate and eluted with the TBS- DTT solution. The number of phages from the selection and control reactions were titered to follow the enrichment of the phage particles from the selection reaction. After each round of selection, 10 -20 colonies were picked from the LB-agar plate of the selection reaction to prepare phagemid DNA. The DNA was sequenced to see if the RBR library was converged to a consensus sequence. The library phagemid from the previous round of selection was used for the next round of phage preparation. The concentration of xUba1 and xUbcH7 enzymes and biotin-xUB was decreased in each round of selection reactions. For the fifth round of selection, 0.06 μM xUba1(UFD), 5 μM xUbcH7, and 0.1 μM biotin–xUB were used, and the reaction time was shortened to 10 min.

### Assaying the activity of the selected RBR mutants of Parkin

The RBR mutants from phage selection were cloned into the pGEX vector using primers GC6 and GC7 between restriction sites BamHI and NotI to express the proteins with a N-terminal GST tag. Ubiquitination of the RBR was set up with 5 μM RBR mutant, 0.1 μM xUba1, 2 μM xUbcH7, and 5 μM HA-xUB in buffer with 50 mM Tris-HCl (pH 7.5), 10 mM MgCl_2_, and 1.5 mM ATP. The reaction was incubated at 37°C for 10 min and was subject to SDS-PAGE and western blotting with an anti-Flag antibody to identify RBR mutants that can mediate the transfer of xUB. To incorporate mutations of RBR into the full-length Parkin gene, primer set GC8 and GC7 was used to amplify the RBR gene. The fragment was assembled with the PCR fragment generated with the GC9/GC10 pair and cloned into the pGEX vector between restriction sites BamHI and NotI for expression of full-length Parkin with RBR mutations. To test the activity of Parkin ubiquitinating Miro1, reactions were set with 20 µM UB, 1 µM Uba1, 10 µM UbcH7, 5 µM Parkin, 5 µM Miro1, and 10 µL 5× reaction buffer (10 mM DTT, 25 mM ATP, 25 mM MgCl_2_, pH 8.0 in PBS) in a total volume of 50 µL. The reaction tubes were shaken at 37℃ overnight before analysis by SDS-PAGE and western blot.

### Proteins expression from Recombinant pET plasmids and pGEX plasmids

Recombinant pET plasmids for the expression of Uba1, UbcH7, UB, and Rab1a, Rab5a, Rab5c, Rab7a, Rab8a, Rab10, Rab13, and CDK5 were transformed into BL21 cells and cultured in 2×YT broth with antibiotics under 37 °C until the OD values of the cultures were in the range of 0.6–0.8. IPTG was added to the cell culture to reach the final concentration of 1 mM to induce protein expression, and the cell culture was incubated overnight under 20°C with agitation at 220 round per minute (rpm) before the cells were harvested by centrifugation (5,500 rpm, 4 °C, 20 min). Cells were resuspended in 20 mL of lysis buffer (50 mM NaH_2_PO_4_, 300 mM NaCl, 10 mM imidazole, pH 8.0) with the addition of 40 mg of lysozyme and 1 mM PMSF, and the suspension was incubated on ice for 30 min. The cells were lysed by sonication on ice and the cell lysates were centrifuged (10,000 rpm, 4°C, 30 min). After centrifugation, the supernatant of the lysate was collected to bind with Ni-NTA beads overnight at 4 °C. The protein was purified by a gravity-flow column with washes by 20 mL of lysis buffer (50 mM NaH_2_PO_4_, 300 mM NaCl, 5 mM imidazole, pH 8.0) once and 20 mL wash buffer (50 mM NaH_2_PO_4_, 300 mM NaCl, 20 mM imidazole, pH 8.0) twice followed by elution with 5 mL of elution buffer (50 mM NaH_2_PO_4_, 300 mM NaCl, 250 mM imidazole, pH 8.0). The eluted protein solution was further dialyzed against a dialysis buffer (50 mM Tris, 50 mM NaCl, 1 mM DTT, pH 8.0) and concentrated.

Recombinant pGEX plasmids for the expression of wt full length Parkin, full length xParkin, wild-type Parkin RBR domain and xParkin RBR mutants were also transformed into BL21 cells and the colonies acquired were used to inoculate 2×YT broth with ampicillin (100 µg/mL) and cultured at 37°C. 1 mM IPTG was added to the cell culture to induce protein expression, and the culture was incubated overnight at 20°C with agitation (220 rpm) before the cells were harvested by centrifugation (5,500 rpm, 4 °C, 20 min). Cells were resuspended in 20 mL of lysis buffer (50 mM Tris, 150 mM NaCl, pH 8.0, freshly added with 1 mg/mL lysozyme, 50 µM ZnCl_2_, 1 mM PMSF, 1 mM proteasome inhibitor, and 10 mM DTT), and incubated on ice for at least 30 min. The cell suspension was sonicated for 30 min on ice and the crude cell lysate was centrifuged at 14,000 rpm for 30 min to remove cell debris. The clear supernatant was collected into a clean 50 mL tube. 1 mL of GST beads was washed 3 times with 1 mL lysis buffer and then added into cell lysate. The mixture was put on a rotator at 4℃ for overnight binding. The next morning, the mixture was applied to a clean affinity column, and was washed sequentially by 15 mL lysis buffer, 2× 15 mL wash buffer (50 mM Tris, 150 mM NaCl, pH 8.0), and eluted with 5 mL elution buffer (50 mM Tris, 150 mM NaCl, 10 mM reduced glutathione, pH 8.0). Eluate was packaged into a dialysis bag and then put into 1 L dialysis buffer (25 mM Tris base, 150 mM NaCl, 0.5 mM DTT, pH 8.0) at 4℃ overnight. The third day, eluate was dialysis for another 4 hours in 1 L of fresh dialysis buffer. Finally, eluate was collected into a 5 kDa-cutoff concentrator tube, centrifuged at 4,000 rpm till the volume was concentrated to 0.5 ∼ 1 mL. The final protein solution was aliquoted and stored at –80℃.

### Tandem purification of HBT-xUB-conjugated proteins from cells expressing the Parkin OUT cascade

pLenti-xParkin and plenti-xParkin with the inactivating C431A mutation were co-transfected with pLenti-HBT-xUB into the HEK293 cells with stable expression of xUba7-xUbcH7 pair ^47^ to express the Parkin OUT cascade and the control OUT cascade in the cell. For transfection, xUba1-xUbcH7 cells were cultured in four 75 cm^2^-flasks and 5 µg of each of pLenti-HBT-xUB and pLenti-myc-xParkin or xParkin C431A plasmid were co-transfected into each flask of cells by DharmaFECT kb DNA transfection reagent (T-2006-01). After 48-hour of transfection, cells were treated with 10 µM MG132 for 4 hours at 37℃ and washed twice with 2 mL ice-cold PBS. The buffer was then removed and cells in each flask of cells were lysed with 1 mL buffer A (8 M urea, 300mM NaCl, 50mM Tris, 50mM NaH_2_PO_4_, 0.5% NP-40, 1 mM PMSF and 125 U/ml benzonase, pH 8.0). The lysate was combined into a clean 15 mL Falcon tube and was sonicated for 2 min with a 5-s on and 5-s off cycle. After sonication, the lysate was centrifuged at 13,000 rpm for 30 min at room temperature. Clear supernatant was collected and total protein concentration was measured by the Branford assay. Next, 35 µL PBS-washed Ni Sepharose™ High Performance (GE healthcare, Cat# 17-5268-01) beads were added to each 1 mg of total protein (at least 10 mg in total), and a final concentration of 10 mM imidazole was added to the bead mixture for binding overnight on a rotator at room temperature. After binding, the Ni-NTA beads were loaded to a gravity column. Beads were washed sequentially with 20-bead volume of buffer A (pH 8.0), buffer A (pH 6.3), and buffer A (pH 6.3) with 10 mM imidazole, and eluted twice with 5-bead volume of buffer B (8 M Urea, 200 mM NaCl, 50 mM Na_2_HPO_4_, 2% SDS, 10 mM EDTA, 100 mM Tris, 25 mM imidazole, pH 4.3). The pH of combined eluate was adjusted to 8.0, and 10 µL Pierce™ high-capacity streptavidin agarose resin (Thermo Scientific, Cat# 20359) was added to each 1 mg of total protein for overnight binding at room temperature. The streptavidin beads were then washed sequentially with 1.5 mL of buffer C (8 M rea, 200 mM NaCl, 2% SDS, 100 mM Tris, pH 8.0), 1.5 mL buffer D (8 M Urea, 1.2 M NaCl, 0.2% SDS, 100 mM Tris, 10% EtOH, 10% Isopropanol, pH 8.0), and finally resuspend with 1.5 mL buffer E (8 M urea, 100 NH_4_HCO_3_, pH 8.0).

### Sample digestion and LC-MS/MS analysis

Residual buffer E was removed and 200 μL of 50 mM NH_4_HCO_3_ was added to each sample, which were then reduced with dithiothreitol (final concentration 1 mM) for 30 min at 25 °C. This was followed by 30 min of alkylation with 5 mM iodoacetamide in the dark. The samples were then digested with 1 μg of Lysyl-endopeptidase (Wako) at room temperature for 2 hours and further digested overnight with 1:50 (w/w) trypsin (Promega) at room temperature. Resulting peptides were acidified with 25 μL of 10% (v/v) formic acid (FA) and 1% (v/v) triflouroacetic acid (TFA) and desalted with a Sep-Pak C18 column (Waters). For desalting, the Sep-Pak column was washed with 1 mL of methanol and 1 mL of 50% (v/v) acetonitrile (ACN). Equilibration was performed with 2 rounds of 1 mL of 0.1% (v/v) TFA in water. The acidified peptides were then loaded to the column, and the column washed with 2 rounds of 1 mL 0.1% (v/v) TFA. Elution was carried out by 2 rounds of 50% (v/v) ACN (400 μL each) and the resulting peptide eluent dried under vacuum. Liquid chromatography coupled to tandem MS (LC-MS/MS) on an Orbitrap Fusion mass spectrometer (Thermo Fisher Scientific) was performed at the Emory Integrated Proteomics Core (EIPC) according to previously published methods^47^. Collected spectra were searched using Proteome Discoverer 2.0 against human UniProt database (90,300 target sequences). Searching parameters included fully tryptic restriction and a parent ion mass tolerance (±20 parts per million). Methionine oxidation (+15.99492 Da), asparagine and glutamine deamidation (+0.98402 Da), lysine ubiquitination (+114.04293 Da), and protein N-terminal acetylation (+42.03670) were variable modifications (up to three were allowed per peptide); cysteine was assigned a fixed carbamidomethyl modification (+57.021465 Da). Percolator was used to filter the PSMs to a false discovery rate of 1%.

### Bioinformatics analysis

To identify and visualize PARKIN substrates, we generated a volcano plot by calculating the p value from the PSM ratio from xPARKIN OUT compared to its associated xPARKIN catalytic inactive control (Log2[PSM ratio OUT/control] > 1). Protein targets with p < 0.1 and a PSM ratio > 2 were classified as potential Parkin substrates. The enriched targets were used as an input to generate the list of gene ontology in the Biological Process category and disease ontology in the up_kw_disease category using DAVID (https://david.ncifcrf.gov/). A threshold of FDR < 0.05 was selected as the significant terms and the top 15 terms were picked to output the bar plot (ggplot2). Target networks were generated using STRING (https://string-db.org). All heat maps with normalized PSM values were generated using the R pheatmap packages.

### *In vitro* ubiquitination of substrate proteins

Reconstituted ubiquitination reactions were set up in 50 μL of reaction buffer (50 mM Tris, 5 mM MgCl_2_, 5 mM ATP, and 1 mM DTT). Substrate proteins (10 μM) were incubated with 0.5 μM wt Uba1, 2 μM wt UbcH7, 10 μM wt HA-UB and 5 μM wt full-length human Parkin protein at 37 °C for 4 hours. 20 μg/mL PINK1 was also added to the reaction to activate Parkin. Parallel control reactions were also set up with each of the enzyme components excluded from the reaction mixture. The reactions were quenched by boiling the samples in the SDS-PAGE loading dye with DTT for 5 min and analyzed by SDS-PAGE and western blot probed with the substrate-specific antibodies.

### Cell-based assay to verify Parkin-catalyzed ubiquitination of the substrate proteins

HEK293T cells were transfected with varying amounts of pLenti-wtParkin plasmid (0-4 µg) for 14 hrs. The cells were then treated with 0.5 μM MG132 for an additional 12 hours and washed twice with ice-cold PBS, pH 7.4, followed by the addition of 1 mL ice-cold RIPA buffer for incubating with the cells at 4 °C for 10 min. The cells were lysed by repeated aspiration through a 21-gauge needle and the cell lysate was transferred to a 1.5 mL tube. The cell debris in the lysate was pelleted by centrifugation at 13,000 rpm. for 20 min at 4 °C, and the supernatant was transferred to a new tube and precleared by adding 1.0 µg of the appropriate control IgG (mouse or rabbit IgG corresponding to the host species of the primary antibody). 20 µL of suspended protein A/G PLUS-agarose was added to the supernatant, and the incubation was continued for 30 min at 4 °C. After the prebinding step, cell lysate containing 2 mg total protein was transferred to a new tube and 30 µL (i.e., 6 µg) primary antibody specific for the substrate proteins (Rab1a, Rab5a, Rab5c, Rab7a, Rab8a, Rab13, CDK5) were added to bind to target proteins in different batches of lysate. The incubation was continued for 2 hours at 4 °C, and 30 µL of resuspended protein A/G PLUS-agarose was added. The tubes were capped and incubated at 4 °C on a rocking platform overnight. The next day, the agarose beads were pelleted by centrifugation at 350 g for 5 min at 4 °C. The beads were then washed three times, each time with 1.0 mL PBS. After the final wash, the beads were resuspended in 40 µL of 1× Laemmli buffer with β-mercaptoethanol. The samples were boiled for 5 min and analyzed by SDS-PAGE. The western blots of the PAGE gels were probed with an anti-UB antibody.

### Protein degradation assays

To examine the effect of Parkin on the steady-state levels of substrates, HEK293T cells (5 × 10^6^ cells) were transiently transfected with varying amounts of pLenti plasmid of Parkin with the DharmaFECT kb transfection kit. Cells were harvested at 16 hours after transfection, and the amount of substrate proteins in the cell lysate was assayed by immunoblotting with substrate-specific antibodies.

### Substrate ubiquitination under mitophagy stimulation

Transfection of pLenti-Parkin into the HEK293T cells was conducted with the DharmaFECT transfection reagents according to the manufacturer’s protocol. After overnight transfection, the cells were treated with 0.5 μM MG132 for an additional 10-12 h. For assaying the effect of Parkin activation on the ubiquitination of the substrates during mitophagy, the cells were treated with 4 μM antimycin A and 10 μM oligomycin (AO) for 1 hour, and substrates was immunoprecipitated with substrate-specific antibodies to assay their ubiquitination level. Briefly, cells were washed twice with ice-cold PBS, pH 7.4, followed by the addition of 1 mL ice-cold RIPA buffer that was allowed to incubate with the cells at 4 °C for 10 min. The cells were disrupted by repeated aspiration through a 21-gauge needle to induce cell lysis and the cell lysate was transferred to a 1.5 mL tube. The cell debris in the lysate was pelleted by centrifugation at 13,000 rpm for 20 min at 4 °C, and the supernatant was transferred to a new tube and precleared by adding 1.0 μg of the appropriate control IgG (normal mouse or rabbit IgG corresponding to the host species of the primary antibody). 20 μL of suspended Protein A/G PLUS-agarose was added to the supernatant, and the incubation was continued for 30 min at 4 °C. After this, cell lysate containing 2 mg total protein was transferred to a new tube and 30 μL (i.e., 6 μg) primary antibody specific for each substrate was added to bind to substrate proteins in the lysate. The incubation was continued for 1 hour at 4 °C, and 30 μL of resuspended protein A/G PLUS-agarose was added. The tubes were capped and incubated at 4 °C on a rocking platform overnight. The next day, the agarose beads were pelleted by centrifugation at 350 g for 5 min at 4 °C. The beads were then washed three times, each time with 1.0 mL PBS. After the final wash, the beads were resuspended in 40 μL of 1× Laemmli buffer with β-mercaptoethanol. The samples were boiled for 5 min and analyzed by SDS-PAGE. The western blots of the PAGE gels were probed with an anti-UB antibody.

### Cancer proteomic and genomic analyses

The protein-protein and RNA-RNA correlation data of Breast Cancer (BRCA) was downloaded from LinkedOmicsKB (https://kb.linkedomics.org/download, data obtained on May 28th, 2024). The protein levels and RNA levels of PRKN (ENSG00000185345.23), RAB1A (ENSG00000138069.18) and RAB7A (ENSG00000075785.13) were extracted from LinkedOmicsKB files ’BRCA_proteomics_gene_abundance_log2_reference_intensity_normalized_Tumor.txt’ and ’BRCA_RNAseq_gene_RSEM_coding_UQ_1500_log2_Tumor.txt’, respectively. For correlation analysis, data of protein levels and RNA levels was merged on patient ID, and 88 samples that were aligned with both proteomic and genomic data were selected for the statistical analysis. Spearman correlation coefficient (rho) and associated p-values were calculated with SciPy package written in Python. Scatter plots and linear regression were drawn with Seaborn package written in Python.

## Funding

This work was supported by the National Institute of Health (R01GM104498 to J.Y. and H.K., R21NS116760 to J.Y. and A.M.M., and R35GM139382 to I.I.), National Science Foundation CAREER Award (2047700) to A.M.M. and National Science Foundation grant 2109051 to J.Y. and MCB-2027902 to I.I. and Natural Science Foundation of China (31971187 to B.Z.) and the Science and Technology Commission of Shanghai Municipality Project (20JC1411200 to B.Z.). L.Z. was supported by a graduate fellowship from the Center for Diagnostics and Therapeutics of Georgia State University. We thank the help of Chunli Yan with the preparation of the method section of the manuscript.

## Author contributions

J.Y., B.Z., and H.K. conceived the project. S.F. performed the proteomics screen of the Parkin substrates. L.Z., X.W., I.H.J., S.E.J., J.Z., and G.H.J. verified Parkin substrates by ubiquitination assays in vitro and in the cell. G.C. engineered Parkin RBR to generate the OUT cascade. B.R.K. and I.I. generated the structural model of Parkin RBR in complex with UB∼UbcH5b. W.W. and A.M.M. performed bioinformatic analysis of Parkin substrates. J.Y., B.Z., H.K., A.M.M., S.F., L.Z., G.C., and W.W. prepared the manuscript with input from all the authors. All authors analyzed and interpreted the results and commented on the manuscript.

## Competing interests

The authors declare that they have no competing interests.

## Data and materials availability

All data needed to evaluate the conclusions in the paper are present in the paper and/or the Supplementary Materials. The proteomics datasets from this study will be deposited to the ProteomeXchange Consortium via the PRIDE partner repository^129^.

---

**Supplemental Figure S1.**
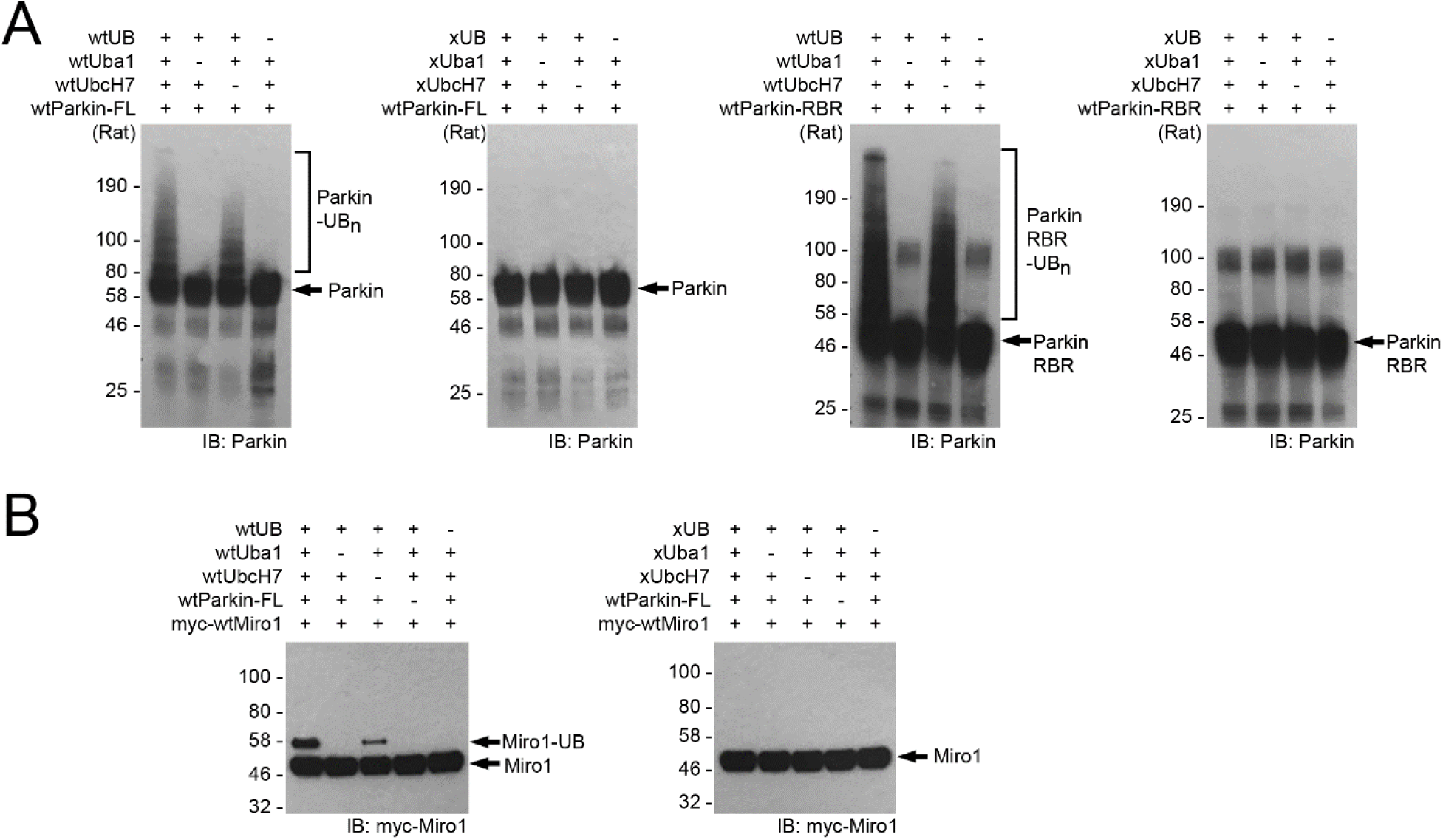
wt Parkin from rat is not active in xUB transfer in the self-ubiquitination reaction and the ubiquitination of Miro1. (A) The full-length rat Parkin and the RBR domain are active in pairing with wt Uba1 (E1) and UbcH7 (E2) for utilizing wt UB for the self-ubiquitination reaction but are incapable of transferring xUB with the xUba1-xUbcH7 pair. (B) The full-length rat Parkin and the RBR domain can transfer wt UB to the Parkin substrate Miro1 but are incapable of transferring xUB to Miro1 with the xUba1-xUbcH7 pair.

**Supplemental Figure S2.**
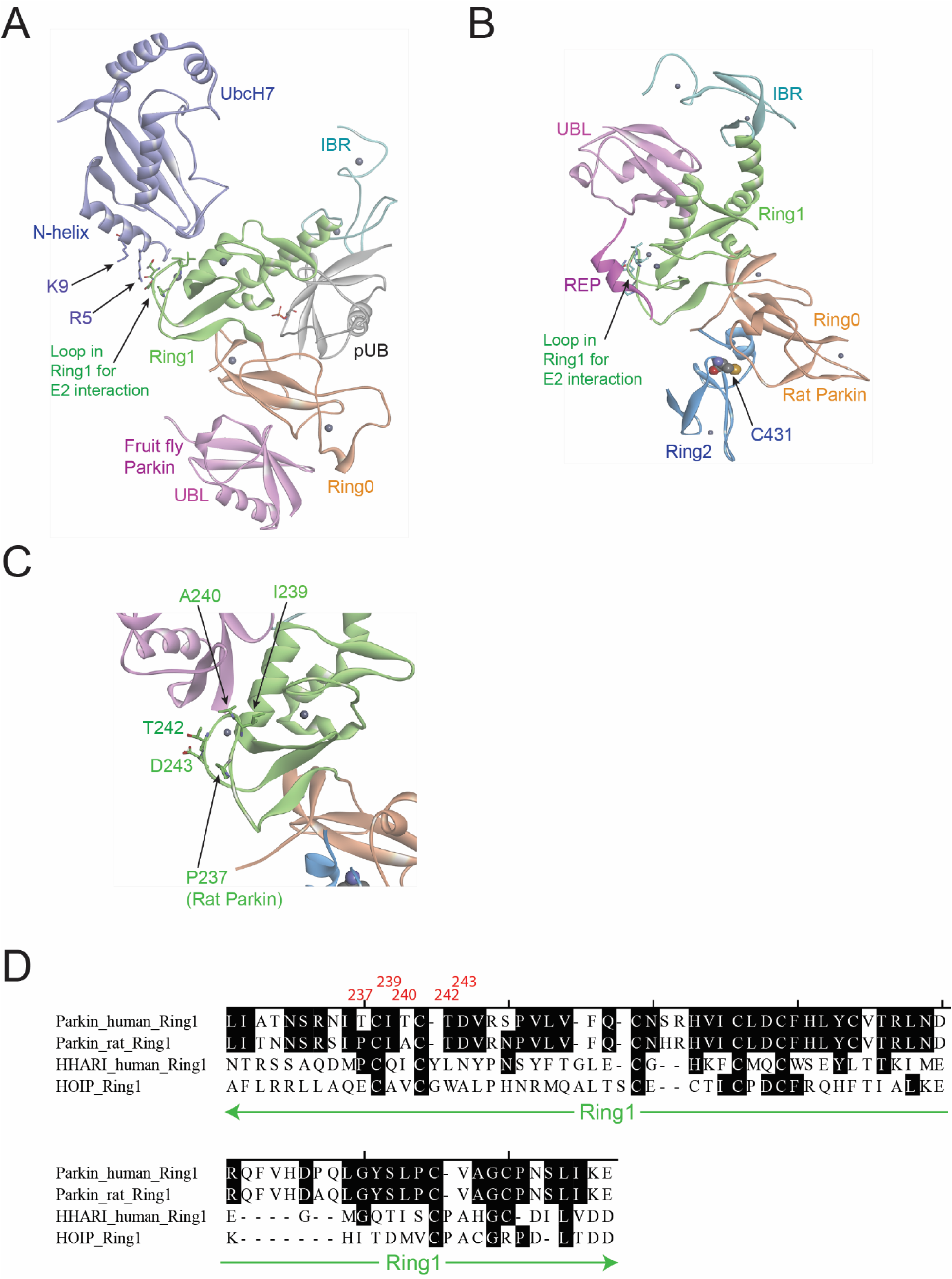
UbcH7-Parkin interaction between the N-terminal helix of UbcH7 and the loop region of the Ring1 domain of Parkin. (A) The crystal structure of Parkin from *B. dorsalis*, an oriental fruit fly, with UbcH7 bound to the Ring1 domain of Parkin (PDB ID: 6DJX)^57^. (B) Crystal structure of rat Parkin showing essential domains of the RBR E3 with the REP element shielding the loop region of the Ring1 domain and blocking the binding of E2 such as UbcH7 to Parkin (PDB ID: 4K95) ^59^. (C) A detailed view of the loop residues in the Ring1 domain of rat Parkin that may bind to the N-terminal helix of UbcH7. The highlighted residues in the loop of Ring1, including P237, I239, A240, T242 and D243, may bind to R5 and K9 residues in the N-terminal helix of UbcH7. (D) The alignment of the protein sequences of the Ring1 domain of RBR E3s Parkin, HHARI and HOIP.

**Supplemental Figure S3.**
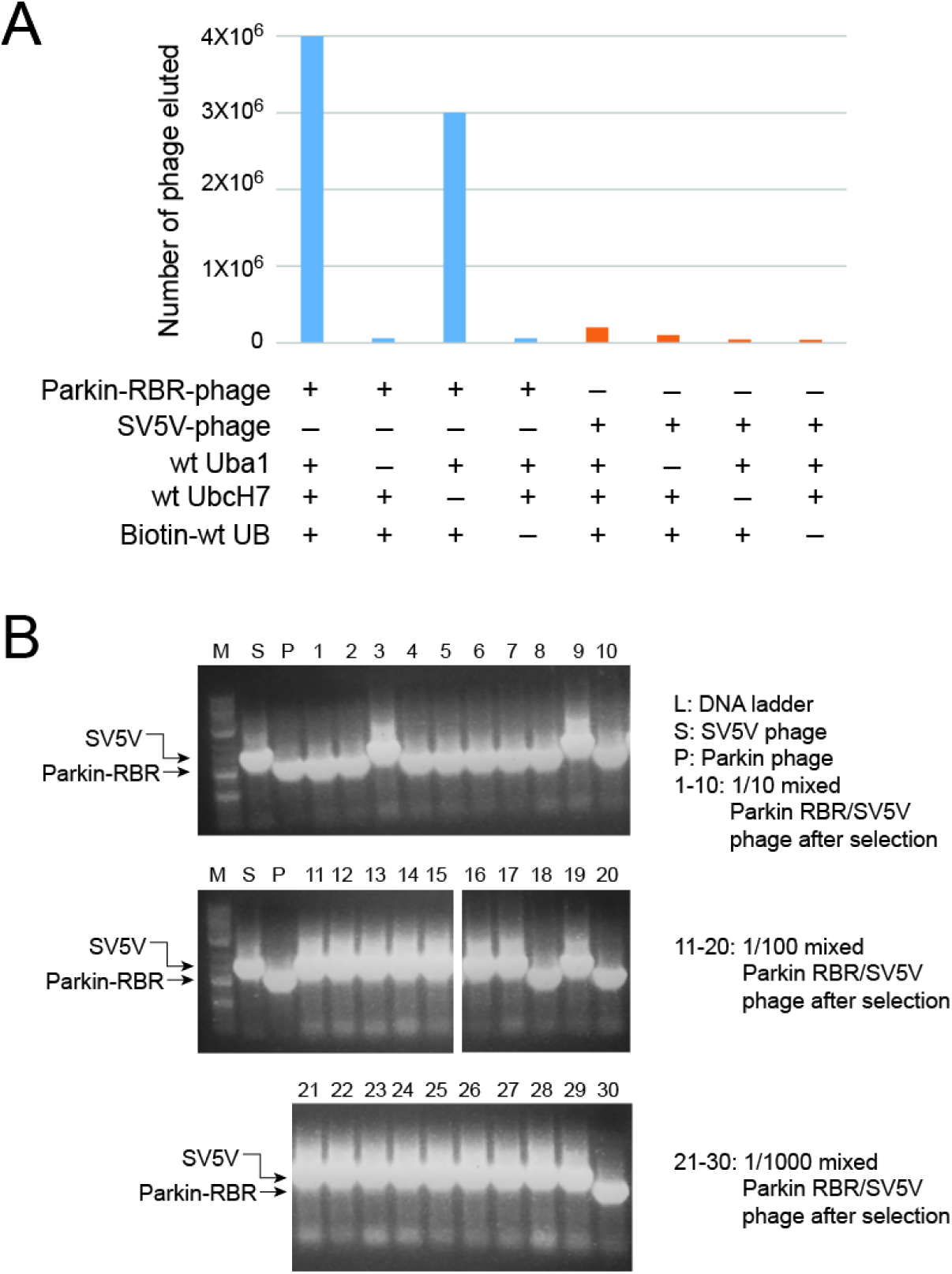
Model selection of the wt Parkin phage displaying the RBR domain of rat Parkin. (A) Phage displaying the wt RBR domain of rat Parkin was reacted with biotin-wt UB and the wt Uba1-UbcH7 pair. In parallel, control reactions were set up with the exclusion of either wt Uba1, UbcH7 or biotin-wt UB from the reaction mixture. In another set of controls, phage displaying a viral protein SV5V with no UB ligase activity were reacted with biotin-wt UB and the wt Uba1-UbcH7 pair. After the reactions, phage particles were bound to the streptavidin plate to retain phage conjugated with biotin-wt UB. The streptavidin plate was then washed to remove unreacted phage, and the phage bound to the plate were titered and their numbers were plotted. (B) Colony PCR reactions to identify phage displaying the wt RBR domain of Parkin after model phage selection with a 1/10, 1/100 and 1/1,000 mixture of RBR displayed Parkin and SV5V displayed Parkin. PCR amplification of the gene encoding the RBR domain of Parkin in the pComb vector in the *E. coli* colonies gave a fragment size of 750 bp compared to the amplification of the SV5V fragment with a size of 900 bp.

**Supplemental Figure S4.**
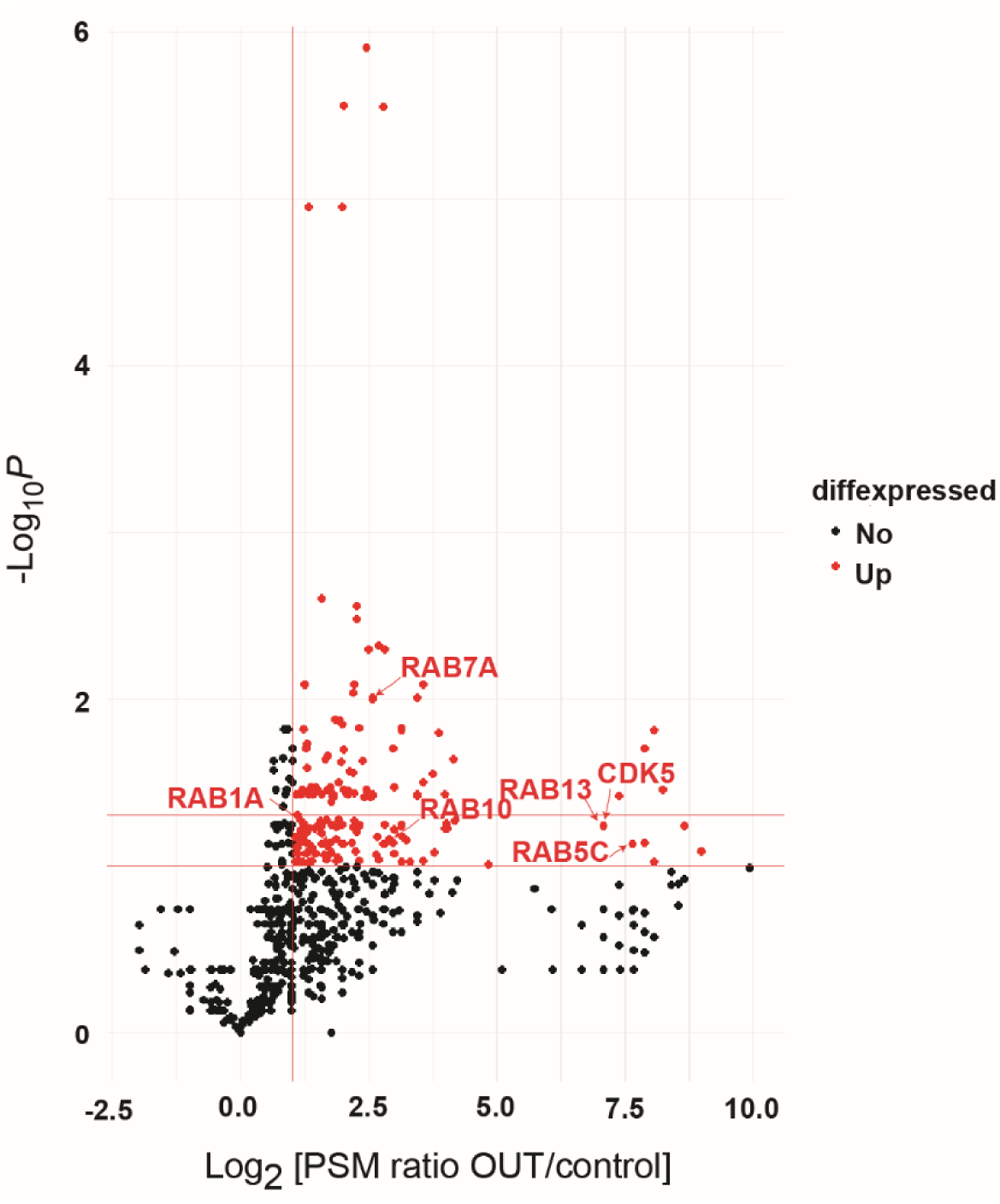
Volcano plot of Parkin substrates identified by the OUT screen. N = 3 independent biological replicates. Red dots designate proteins with Log2[PSM ratio OUT/control] >1 and -Log10*P* > 1.

**Supplemental Figure S5.**
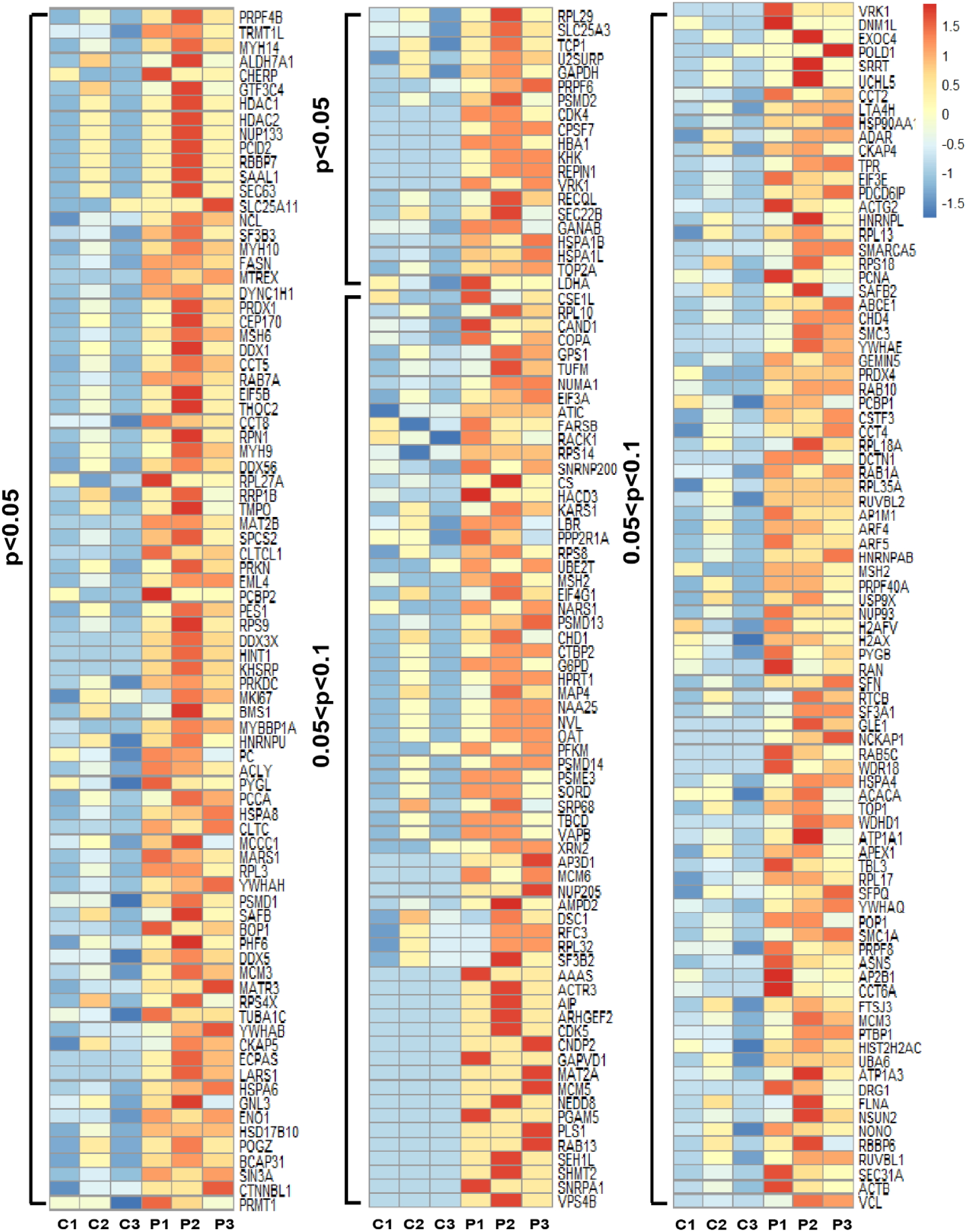
Heat map with normalized PSM values for the total identified 256 protein Parkin substrates with p value as shown in the graph. P1-P3: 3 replicates from OUT cells expressing the functional OUT cascade of Parkin; C1-C3: 3 replicates from the control cells with C431 mutation in xParkin.

**Supplemental Figure S6.**
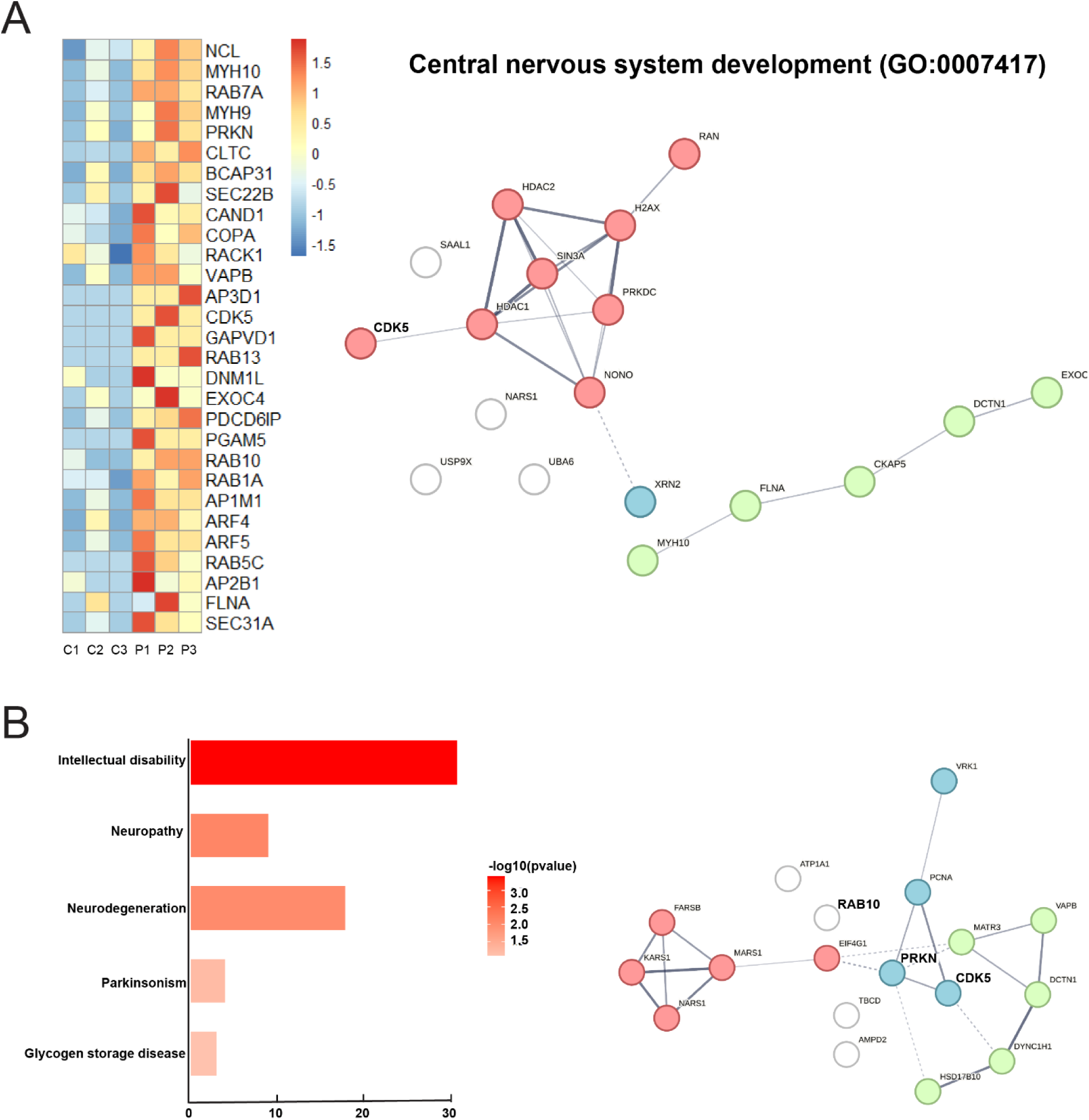
Analysis of Parkin substrates associated with neurodevelopment and diseases. (A) *Left,* Heatmap with normalized PSM values for the targets identified from OUT that were assigned to central nervous system development. *Right,* STRING protein-protein interaction network for targets identified from OUT that were assigned to central nervous system development (GO:0007417). (B) *Left*, Visualization of top significant terms from the list of disease-associated annotations from DAVID. *Right,* STRING protein-protein interaction network for the targets involved in neurodegeneration (KW-0523).

**Supplemental Figure S7.**
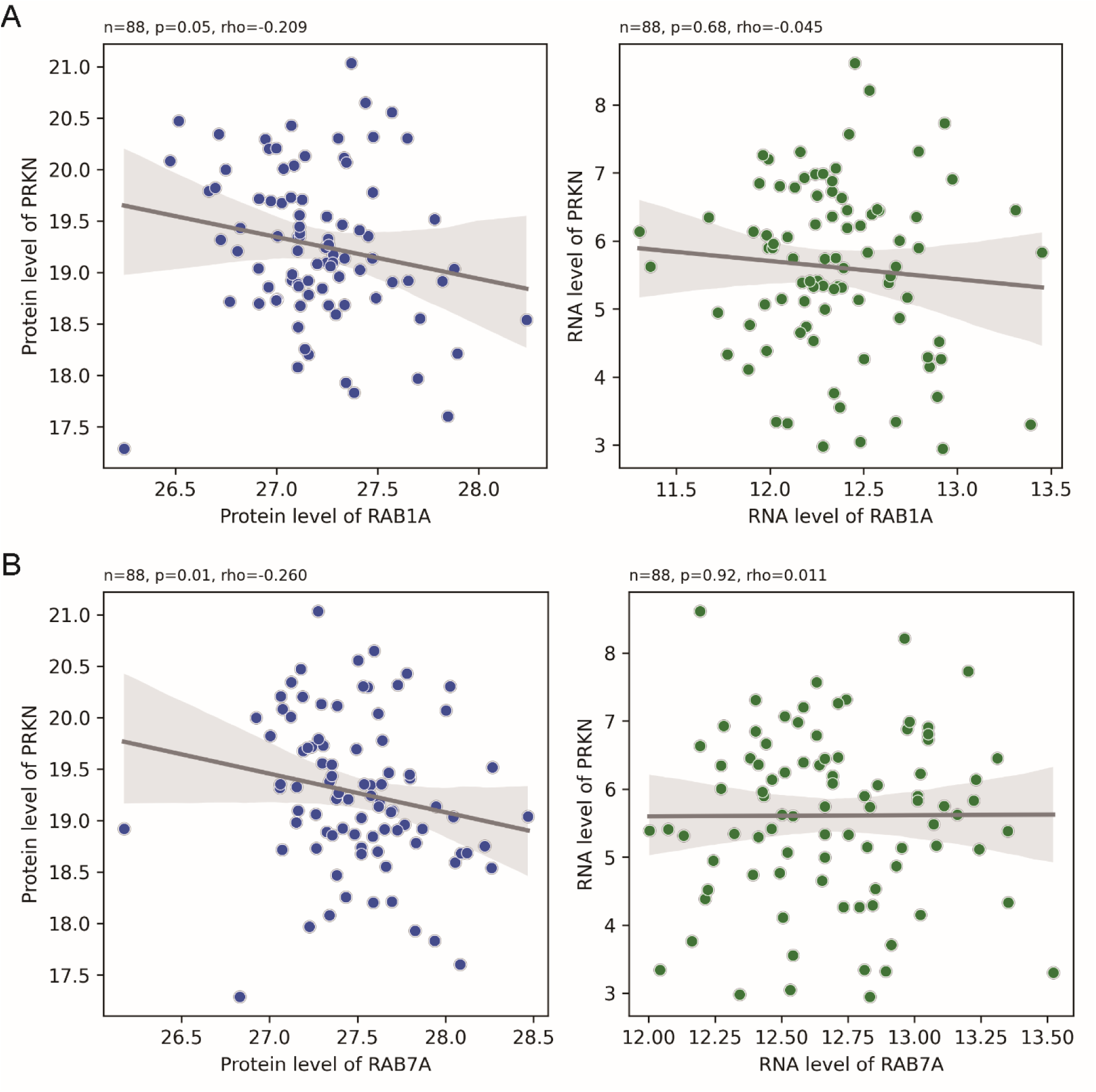
The protein level of Parkin is reciprocally correlated with those of Rab1a and Rab7a in human breast cancer tissues, whereas there are no significant correlations between the mRNA levels of Parkin and those of the Rab proteins. Data from each sample of breast cancer tissues were obtained from the LinkedOmicsKB proteogenomics database (https://kb.linkedomics.org/) and analyzed by Spearman correlation coefficient (rho) with associated p-values calculated with SciPy package written in Python.

## Notes

### Competing Interest Statement

The authors have declared no competing interest.

## References

1. Dove, K. K.; Klevit, R. E., RING-Between-RING E3 Ligases: Emerging Themes amid the Variations. J Mol Biol 2017, 429 (22), 3363–3375. https://www.ncbi.nlm.nih.gov/pubmed/28827147.

2. Trempe, J. F.; Gehring, K., Structural Mechanisms of Mitochondrial Quality Control Mediated by PINK1 and Parkin. J Mol Biol 2023, 435 (12), 168090. https://www.ncbi.nlm.nih.gov/pubmed/37054910.

3. Yamano, K.; Matsuda, N.; Tanaka, K., The ubiquitin signal and autophagy: an orchestrated dance leading to mitochondrial degradation. EMBO Rep 2016, 17 (3), 300–16. https://www.ncbi.nlm.nih.gov/pubmed/26882551.

4. Pickrell, A. M.; Youle, R. J., The roles of PINK1, parkin, and mitochondrial fidelity in Parkinson’s disease. Neuron 2015, 85 (2), 257–73. https://www.ncbi.nlm.nih.gov/pubmed/25611507.

5. Bernardini, J. P.; Lazarou, M.; Dewson, G., Parkin and mitophagy in cancer. Oncogene 2017, 36 (10), 1315–1327. https://www.ncbi.nlm.nih.gov/pubmed/27593930.

6. Poole, L. P.; Macleod, K. F., Mitophagy in tumorigenesis and metastasis. Cell Mol Life Sci 2021, 78 (8), 3817–3851. https://www.ncbi.nlm.nih.gov/pubmed/33580835.

7. Lazarou, M.; Sliter, D. A.; Kane, L. A.; Sarraf, S. A.; Wang, C.; Burman, J. L.; Sideris, D. P.; Fogel, A. I.; Youle, R. J., The ubiquitin kinase PINK1 recruits autophagy receptors to induce mitophagy. Nature 2015, 524 (7565), 309–14. https://www.ncbi.nlm.nih.gov/pubmed/26266977.

8. Harper, J. W.; Ordureau, A.; Heo, J. M., Building and decoding ubiquitin chains for mitophagy. Nat Rev Mol Cell Biol 2018, 19 (2), 93–108. https://www.ncbi.nlm.nih.gov/pubmed/29358684.

9. Ordureau, A.; Sarraf, S. A.; Duda, D. M.; Heo, J. M.; Jedrychowski, M. P.; Sviderskiy, V. O.; Olszewski, J. L.; Koerber, J. T.; Xie, T.; Beausoleil, S. A.; Wells, J. A.; Gygi, S. P.; Schulman, B. A.; Harper, J. W., Quantitative proteomics reveal a feedforward mechanism for mitochondrial PARKIN translocation and ubiquitin chain synthesis. Mol Cell 2014, 56 (3), 360–75. https://www.ncbi.nlm.nih.gov/pubmed/25284222.

10. Narendra, D.; Tanaka, A.; Suen, D. F.; Youle, R. J., Parkin is recruited selectively to impaired mitochondria and promotes their autophagy. J Cell Biol 2008, 183 (5), 795–803. https://www.ncbi.nlm.nih.gov/pubmed/19029340.

11. Kane, L. A.; Lazarou, M.; Fogel, A. I.; Li, Y.; Yamano, K.; Sarraf, S. A.; Banerjee, S.; Youle, R. J., PINK1 phosphorylates ubiquitin to activate Parkin E3 ubiquitin ligase activity. J Cell Biol 2014, 205 (2), 143–53. https://www.ncbi.nlm.nih.gov/pubmed/24751536.

12. Jin, S. M.; Lazarou, M.; Wang, C.; Kane, L. A.; Narendra, D. P.; Youle, R. J., Mitochondrial membrane potential regulates PINK1 import and proteolytic destabilization by PARL. J Cell Biol 2010, 191 (5), 933–42. https://www.ncbi.nlm.nih.gov/pubmed/21115803.

13. Koyano, F.; Okatsu, K.; Kosako, H.; Tamura, Y.; Go, E.; Kimura, M.; Kimura, Y.; Tsuchiya, H.; Yoshihara, H.; Hirokawa, T.; Endo, T.; Fon, E. A.; Trempe, J. F.; Saeki, Y.; Tanaka, K.; Matsuda, N., Ubiquitin is phosphorylated by PINK1 to activate parkin. Nature 2014, 510 (7503), 162–6. https://www.ncbi.nlm.nih.gov/pubmed/24784582.

14. Panicker, N.; Ge, P.; Dawson, V. L.; Dawson, T. M., The cell biology of Parkinson’s disease. J Cell Biol 2021, 220 (4). https://www.ncbi.nlm.nih.gov/pubmed/33749710.

15. Kasten, M.; Hartmann, C.; Hampf, J.; Schaake, S.; Westenberger, A.; Vollstedt, E. J.; Balck, A.; Domingo, A.; Vulinovic, F.; Dulovic, M.; Zorn, I.; Madoev, H.; Zehnle, H.; Lembeck, C. M.; Schawe, L.; Reginold, J.; Huang, J.; Konig, I. R.; Bertram, L.; Marras, C.; Lohmann, K.; Lill, C. M.; Klein, C., Genotype-Phenotype Relations for the Parkinson’s Disease Genes Parkin, PINK1, DJ1: MDSGene Systematic Review. *Mov Disord* 2018, *33* (5), 730-741. https://www.ncbi.nlm.nih.gov/pubmed/29644727.

16. Van Laar, V. S.; Arnold, B.; Cassady, S. J.; Chu, C. T.; Burton, E. A.; Berman, S. B., Bioenergetics of neurons inhibit the translocation response of Parkin following rapid mitochondrial depolarization. Hum Mol Genet 2011, 20 (5), 927–40. https://www.ncbi.nlm.nih.gov/pubmed/21147754.

17. Lee, J. J.; Sanchez-Martinez, A.; Martinez Zarate, A.; Beninca, C.; Mayor, U.; Clague, M. J.; Whitworth, A. J., Basal mitophagy is widespread in Drosophila but minimally affected by loss of Pink1 or parkin. J Cell Biol 2018, 217 (5), 1613–1622. https://www.ncbi.nlm.nih.gov/pubmed/29500189.

18. McWilliams, T. G.; Barini, E.; Pohjolan-Pirhonen, R.; Brooks, S. P.; Singh, F.; Burel, S.; Balk, K.; Kumar, A.; Montava-Garriga, L.; Prescott, A. R.; Hassoun, S. M.; Mouton-Liger, F.; Ball, G.; Hills, R.; Knebel, A.; Ulusoy, A.; Di Monte, D. A.; Tamjar, J.; Antico, O.; Fears, K.; Smith, L.; Brambilla, R.; Palin, E.; Valori, M.; Eerola-Rautio, J.; Tienari, P.; Corti, O.; Dunnett, S. B.; Ganley, I. G.; Suomalainen, A.; Muqit, M. M. K., Phosphorylation of Parkin at serine 65 is essential for its activation in vivo. Open Biol 2018, 8 (11). https://www.ncbi.nlm.nih.gov/pubmed/30404819.

19. McWilliams, T. G.; Prescott, A. R.; Montava-Garriga, L.; Ball, G.; Singh, F.; Barini, E.; Muqit, M. M. K.; Brooks, S. P.; Ganley, I. G., Basal Mitophagy Occurs Independently of PINK1 in Mouse Tissues of High Metabolic Demand. Cell Metab 2018, 27 (2), 439–449 e5. https://www.ncbi.nlm.nih.gov/pubmed/29337137.

20. Gegg, M. E.; Cooper, J. M.; Chau, K. Y.; Rojo, M.; Schapira, A. H.; Taanman, J. W., Mitofusin 1 and mitofusin 2 are ubiquitinated in a PINK1/parkin-dependent manner upon induction of mitophagy. Hum Mol Genet 2010, 19 (24), 4861–70. https://www.ncbi.nlm.nih.gov/pubmed/20871098.

21. Tanaka, A.; Cleland, M. M.; Xu, S.; Narendra, D. P.; Suen, D. F.; Karbowski, M.; Youle, R. J., Proteasome and p97 mediate mitophagy and degradation of mitofusins induced by Parkin. J Cell Biol 2010, 191 (7), 1367–80. https://www.ncbi.nlm.nih.gov/pubmed/21173115.

22. Wang, X.; Winter, D.; Ashrafi, G.; Schlehe, J.; Wong, Y. L.; Selkoe, D.; Rice, S.; Steen, J.; LaVoie, M. J.; Schwarz, T. L., PINK1 and Parkin target Miro for phosphorylation and degradation to arrest mitochondrial motility. Cell 2011, 147 (4), 893–906. https://www.ncbi.nlm.nih.gov/pubmed/22078885.

23. Safiulina, D.; Kuum, M.; Choubey, V.; Gogichaishvili, N.; Liiv, J.; Hickey, M. A.; Cagalinec, M.; Mandel, M.; Zeb, A.; Liiv, M.; Kaasik, A., Miro proteins prime mitochondria for Parkin translocation and mitophagy. Embo J 2019, 38 (2). https://www.ncbi.nlm.nih.gov/pubmed/30504269.

24. Shin, J. H.; Ko, H. S.; Kang, H.; Lee, Y.; Lee, Y. I.; Pletinkova, O.; Troconso, J. C.; Dawson, V. L.; Dawson, T. M., PARIS (ZNF746) repression of PGC-1alpha contributes to neurodegeneration in Parkinson’s disease. Cell 2011, 144 (5), 689–702. https://www.ncbi.nlm.nih.gov/pubmed/21376232.

25. Siddiqui, A.; Rane, A.; Rajagopalan, S.; Chinta, S. J.; Andersen, J. K., Detrimental effects of oxidative losses in parkin activity in a model of sporadic Parkinson’s disease are attenuated by restoration of PGC1alpha. Neurobiol Dis 2016, 93, 115–20. https://www.ncbi.nlm.nih.gov/pubmed/27185595.

26. Stevens, D. A.; Lee, Y.; Kang, H. C.; Lee, B. D.; Lee, Y. I.; Bower, A.; Jiang, H.; Kang, S. U.; Andrabi, S. A.; Dawson, V. L.; Shin, J. H.; Dawson, T. M., Parkin loss leads to PARIS-dependent declines in mitochondrial mass and respiration. Proc Natl Acad Sci U S A 2015, 112 (37), 11696–701. https://www.ncbi.nlm.nih.gov/pubmed/26324925.

27. Vandiver, M. S.; Paul, B. D.; Xu, R.; Karuppagounder, S.; Rao, F.; Snowman, A. M.; Ko, H. S.; Lee, Y. I.; Dawson, V. L.; Dawson, T. M.; Sen, N.; Snyder, S. H., Sulfhydration mediates neuroprotective actions of parkin. Nat Commun 2013, 4, 1626. https://www.ncbi.nlm.nih.gov/pubmed/23535647.

28. Ozawa, K.; Komatsubara, A. T.; Nishimura, Y.; Sawada, T.; Kawafune, H.; Tsumoto, H.; Tsuji, Y.; Zhao, J.; Kyotani, Y.; Tanaka, T.; Takahashi, R.; Yoshizumi, M., S-nitrosylation regulates mitochondrial quality control via activation of parkin. Sci Rep 2013, 3, 2202. https://www.ncbi.nlm.nih.gov/pubmed/23857542.

29. Lee, Y.; Stevens, D. A.; Kang, S. U.; Jiang, H.; Lee, Y. I.; Ko, H. S.; Scarffe, L. A.; Umanah, G. E.; Kang, H.; Ham, S.; Kam, T. I.; Allen, K.; Brahmachari, S.; Kim, J. W.; Neifert, S.; Yun, S. P.; Fiesel, F. C.; Springer, W.; Dawson, V. L.; Shin, J. H.; Dawson, T. M., PINK1 Primes Parkin-Mediated Ubiquitination of PARIS in Dopaminergic Neuronal Survival. Cell Rep 2017, 18 (4), 918–932. https://www.ncbi.nlm.nih.gov/pubmed/28122242.

30. Lee, S. B.; Kim, J. J.; Nam, H. J.; Gao, B.; Yin, P.; Qin, B.; Yi, S. Y.; Ham, H.; Evans, D.; Kim, S. H.; Zhang, J.; Deng, M.; Liu, T.; Zhang, H.; Billadeau, D. D.; Wang, L.; Giaime, E.; Shen, J.; Pang, Y. P.; Jen, J.; van Deursen, J. M.; Lou, Z., Parkin Regulates Mitosis and Genomic Stability through Cdc20/Cdh1. Mol Cell 2015, 60 (1), 21–34. https://www.ncbi.nlm.nih.gov/pubmed/26387737.

31. Johnson, B. N.; Berger, A. K.; Cortese, G. P.; Lavoie, M. J., The ubiquitin E3 ligase parkin regulates the proapoptotic function of Bax. Proc Natl Acad Sci U S A 2012, 109 (16), 6283–8. https://www.ncbi.nlm.nih.gov/pubmed/22460798.

32. Bernardini, J. P.; Brouwer, J. M.; Tan, I. K.; Sandow, J. J.; Huang, S.; Stafford, C. A.; Bankovacki, A.; Riffkin, C. D.; Wardak, A. Z.; Czabotar, P. E.; Lazarou, M.; Dewson, G., Parkin inhibits BAK and BAX apoptotic function by distinct mechanisms during mitophagy. Embo J 2019, 38 (2). https://www.ncbi.nlm.nih.gov/pubmed/30573668.

33. Liu, K.; Li, F.; Han, H.; Chen, Y.; Mao, Z.; Luo, J.; Zhao, Y.; Zheng, B.; Gu, W.; Zhao, W., Parkin Regulates the Activity of Pyruvate Kinase M2. J Biol Chem 2016, 291 (19), 10307–17. https://www.ncbi.nlm.nih.gov/pubmed/26975375.

34. Perwez, A.; Wahabi, K.; Rizvi, M. A., Parkin: A targetable linchpin in human malignancies. Biochim Biophys Acta Rev Cancer 2021, 1876 (1), 188533. https://www.ncbi.nlm.nih.gov/pubmed/33785381.

35. Mengesdorf, T.; Jensen, P. H.; Mies, G.; Aufenberg, C.; Paschen, W., Down-regulation of parkin protein in transient focal cerebral ischemia: A link between stroke and degenerative disease? Proc Natl Acad Sci U S A 2002, 99 (23), 15042–7. https://www.ncbi.nlm.nih.gov/pubmed/12415119.

36. Mira, M. T.; Alcais, A.; Nguyen, V. T.; Moraes, M. O.; Di Flumeri, C.; Vu, H. T.; Mai, C. P.; Nguyen, T. H.; Nguyen, N. B.; Pham, X. K.; Sarno, E. N.; Alter, A.; Montpetit, A.; Moraes, M. E.; Moraes, J. R.; Dore, C.; Gallant, C. J.; Lepage, P.; Verner, A.; Van De Vosse, E.; Hudson, T. J.; Abel, L.; Schurr, E., Susceptibility to leprosy is associated with PARK2 and PACRG. Nature 2004, 427 (6975), 636–40. https://www.ncbi.nlm.nih.gov/pubmed/14737177.

37. Bayne, A. N.; Trempe, J. F., Mechanisms of PINK1, ubiquitin and Parkin interactions in mitochondrial quality control and beyond. Cell Mol Life Sci 2019, 76 (23), 4589–4611. https://www.ncbi.nlm.nih.gov/pubmed/31254044.

38. Yin, C. L.; Chen, H. I.; Li, L. H.; Chien, Y. L.; Liao, H. M.; Chou, M. C.; Chou, W. J.; Tsai, W. C.; Chiu, Y. N.; Wu, Y. Y.; Lo, C. Z.; Wu, J. Y.; Chen, Y. T.; Gau, S. S., Genome-wide analysis of copy number variations identifies PARK2 as a candidate gene for autism spectrum disorder. Mol Autism 2016, 7, 23. https://www.ncbi.nlm.nih.gov/pubmed/27042285.

39. Scheuerle, A.; Wilson, K., PARK2 copy number aberrations in two children presenting with autism spectrum disorder: further support of an association and possible evidence for a new microdeletion/microduplication syndrome. Am J Med Genet B Neuropsychiatr Genet 2011, *156B* (4), 413–20. https://www.ncbi.nlm.nih.gov/pubmed/21360662.

40. Deshaies, R. J.; Joazeiro, C. A., RING domain E3 ubiquitin ligases. Annu Rev Biochem 2009, 78, 399–434. http://www.ncbi.nlm.nih.gov/entrez/query.fcgi?cmd=Retrieve&db=PubMed&dopt=Citation&list_uids=19489725

41. George, A. J.; Hoffiz, Y. C.; Charles, A. J.; Zhu, Y.; Mabb, A. M., A Comprehensive Atlas of E3 Ubiquitin Ligase Mutations in Neurological Disorders. Front Genet 2018, 9, 29. https://www.ncbi.nlm.nih.gov/pubmed/29491882.

42. Marin, I.; Lucas, J. I.; Gradilla, A. C.; Ferrus, A., Parkin and relatives: the RBR family of ubiquitin ligases. Physiol Genomics 2004, 17 (3), 253–63. https://www.ncbi.nlm.nih.gov/pubmed/15152079.

43. Ye, Y.; Rape, M., Building ubiquitin chains: E2 enzymes at work. Nat Rev Mol Cell Biol 2009, 10 (11), 755–64. http://www.ncbi.nlm.nih.gov/entrez/query.fcgi?cmd=Retrieve&db=PubMed&dopt=Citation&list_uids=19851334

44. Wenzel, D. M.; Stoll, K. E.; Klevit, R. E., E2s: structurally economical and functionally replete. Biochem J 2011, 433 (1), 31–42. http://www.ncbi.nlm.nih.gov/pubmed/21158740.

45. Jin, J.; Li, X.; Gygi, S. P.; Harper, J. W., Dual E1 activation systems for ubiquitin differentially regulate E2 enzyme charging. Nature 2007, 447 (7148), 1135–8. http://www.ncbi.nlm.nih.gov/entrez/query.fcgi?cmd=Retrieve&db=PubMed&dopt=Citation&list_uids=17597759

46. Zhao, B.; Tsai, Y. C.; Jin, B.; Wang, B.; Wang, Y.; Zhou, H.; Carpenter, T.; Weissman, A. M.; Yin, J., Protein Engineering in the Ubiquitin System: Tools for Discovery and Beyond. Pharmacol Rev 2020, 72 (2), 380–413. https://www.ncbi.nlm.nih.gov/pubmed/32107274.

47. Wang, Y.; Liu, X.; Zhou, L.; Duong, D.; Bhuripanyo, K.; Zhao, B.; Zhou, H.; Liu, R.; Bi, Y.; Kiyokawa, H.; Yin, J., Identifying the ubiquitination targets of E6AP by orthogonal ubiquitin transfer. Nat Commun 2017, 8 (1), 2232. https://www.ncbi.nlm.nih.gov/pubmed/29263404.

48. Bhuripanyo, K.; Wang, Y.; Liu, X.; Zhou, L.; Liu, R.; Duong, D.; Zhao, B.; Bi, Y.; Zhou, H.; Chen, G.; Seyfried, N. T.; Chazin, W. J.; Kiyokawa, H.; Yin, J., Identifying the substrate proteins of U-box E3s E4B and CHIP by orthogonal ubiquitin transfer. Sci Adv 2018, 4 (1), e1701393. https://www.ncbi.nlm.nih.gov/pubmed/29326975.

49. Wang, Y.; Liu, R.; Liao, J.; Jiang, L.; Jeong, G. H.; Zhou, L.; Polite, M.; Duong, D.; Seyfried, N. T.; Wang, H.; Kiyokawa, H.; Yin, J., Orthogonal ubiquitin transfer reveals human papillomavirus E6 downregulates nuclear transport to disarm interferon-gamma dependent apoptosis of cervical cancer cells. FASEB J 2021, 35 (11), e21986. https://www.ncbi.nlm.nih.gov/pubmed/34662469.

50. Wang, Y.; Fang, S.; Chen, G.; Ganti, R.; Chernova, T. A.; Zhou, L.; Duong, D.; Kiyokawa, H.; Li, M.; Zhao, B.; Shcherbik, N.; Chernoff, Y. O.; Yin, J., Regulation of the endocytosis and prion-chaperoning machineries by yeast E3 ubiquitin ligase Rsp5 as revealed by orthogonal ubiquitin transfer. Cell Chem Biol 2021, 28, 1283–1297. https://www.ncbi.nlm.nih.gov/pubmed/33667410.

51. Martinez, A.; Lectez, B.; Ramirez, J.; Popp, O.; Sutherland, J. D.; Urbe, S.; Dittmar, G.; Clague, M. J.; Mayor, U., Quantitative proteomic analysis of Parkin substrates in Drosophila neurons. Mol Neurodegener 2017, 12 (1), 29. https://www.ncbi.nlm.nih.gov/pubmed/28399880.

52. Watanabe, M.; Saeki, Y.; Takahashi, H.; Ohtake, F.; Yoshida, Y.; Kasuga, Y.; Kondo, T.; Yaguchi, H.; Suzuki, M.; Ishida, H.; Tanaka, K.; Hatakeyama, S., A substrate-trapping strategy to find E3 ubiquitin ligase substrates identifies Parkin and TRIM28 targets. Commun Biol 2020, 3 (1), 592. https://www.ncbi.nlm.nih.gov/pubmed/33082525.

53. Ordureau, A.; Paulo, J. A.; Zhang, J.; An, H.; Swatek, K. N.; Cannon, J. R.; Wan, Q.; Komander, D.; Harper, J. W., Global Landscape and Dynamics of Parkin and USP30-Dependent Ubiquitylomes in iNeurons during Mitophagic Signaling. Mol Cell 2020, 77 (5), 1124–1142 e10. https://www.ncbi.nlm.nih.gov/pubmed/32142685.

54. Antico, O.; Ordureau, A.; Stevens, M.; Singh, F.; Nirujogi, R. S.; Gierlinski, M.; Barini, E.; Rickwood, M. L.; Prescott, A.; Toth, R.; Ganley, I. G.; Harper, J. W.; Muqit, M. M. K., Global ubiquitylation analysis of mitochondria in primary neurons identifies endogenous Parkin targets following activation of PINK1. Sci Adv 2021, 7 (46), eabj0722. https://www.ncbi.nlm.nih.gov/pubmed/34767452.

55. Sarraf, S. A.; Raman, M.; Guarani-Pereira, V.; Sowa, M. E.; Huttlin, E. L.; Gygi, S. P.; Harper, J. W., Landscape of the PARKIN-dependent ubiquitylome in response to mitochondrial depolarization. Nature 2013, 496 (7445), 372–6. http://www.ncbi.nlm.nih.gov/pubmed/23503661.

56. Zhao, B.; Bhuripanyo, K.; Zhang, K.; Kiyokawa, H.; Schindelin, H.; Yin, J., Orthogonal Ubiquitin Transfer through Engineered E1-E2 Cascades for Protein Ubiquitination. Chem. Biol. 2012, 19 (10), 1265–77. https://www.ncbi.nlm.nih.gov/pubmed/23102221.

57. Sauve, V.; Sung, G.; Soya, N.; Kozlov, G.; Blaimschein, N.; Miotto, L. S.; Trempe, J. F.; Lukacs, G. L.; Gehring, K., Mechanism of parkin activation by phosphorylation. Nat Struct Mol Biol 2018, 25 (7), 623–630. https://www.ncbi.nlm.nih.gov/pubmed/29967542.

58. Condos, T. E.; Dunkerley, K. M.; Freeman, E. A.; Barber, K. R.; Aguirre, J. D.; Chaugule, V. K.; Xiao, Y.; Konermann, L.; Walden, H.; Shaw, G. S., Synergistic recruitment of UbcH7∼Ub and phosphorylated Ubl domain triggers parkin activation. Embo J 2018, 37 (23). https://www.ncbi.nlm.nih.gov/pubmed/30446597.

59. Trempe, J. F.; Sauve, V.; Grenier, K.; Seirafi, M.; Tang, M. Y.; Menade, M.; Al-Abdul-Wahid, S.; Krett, J.; Wong, K.; Kozlov, G.; Nagar, B.; Fon, E. A.; Gehring, K., Structure of parkin reveals mechanisms for ubiquitin ligase activation. Science 2013, 340 (6139), 1451–5. http://www.ncbi.nlm.nih.gov/pubmed/23661642.

60. Lechtenberg, B. C.; Rajput, A.; Sanishvili, R.; Dobaczewska, M. K.; Ware, C. F.; Mace, P. D.; Riedl, S. J., Structure of a HOIP/E2∼ubiquitin complex reveals RBR E3 ligase mechanism and regulation. Nature 2016, 529 (7587), 546–50. http://www.ncbi.nlm.nih.gov/pubmed/26789245.

61. Chew, K. C.; Matsuda, N.; Saisho, K.; Lim, G. G.; Chai, C.; Tan, H. M.; Tanaka, K.; Lim, K. L., Parkin mediates apparent E2-independent monoubiquitination in vitro and contains an intrinsic activity that catalyzes polyubiquitination. PLoS One 2011, 6 (5), e19720. https://www.ncbi.nlm.nih.gov/pubmed/21625422.

62. Li, T.; Chen, X.; Garbutt, K. C.; Zhou, P.; Zheng, N., Structure of DDB1 in complex with a paramyxovirus V protein: viral hijack of a propeller cluster in ubiquitin ligase. Cell 2006, 124 (1), 105–17. https://www.ncbi.nlm.nih.gov/pubmed/16413485.

63. Agarwal, E.; Goldman, A. R.; Tang, H. Y.; Kossenkov, A. V.; Ghosh, J. C.; Languino, L. R.; Vaira, V.; Speicher, D. W.; Altieri, D. C., A cancer ubiquitome landscape identifies metabolic reprogramming as target of Parkin tumor suppression. Sci Adv 2021, 7 (35). https://www.ncbi.nlm.nih.gov/pubmed/34433563.

64. Bonet-Ponce, L.; Cookson, M. R., The role of Rab GTPases in the pathobiology of Parkinson’ disease. Curr Opin Cell Biol 2019, 59, 73–80. https://www.ncbi.nlm.nih.gov/pubmed/31054512.

65. Abeliovich, A.; Gitler, A. D., Defects in trafficking bridge Parkinson’s disease pathology and genetics. Nature 2016, 539 (7628), 207–216. https://www.ncbi.nlm.nih.gov/pubmed/27830778.

66. Singh, P. K.; Muqit, M. M. K., Parkinson’s: A Disease of Aberrant Vesicle Trafficking. Annu Rev Cell Dev Biol 2020, 36, 237–264. https://www.ncbi.nlm.nih.gov/pubmed/32749865.

67. Smith, P. D.; Crocker, S. J.; Jackson-Lewis, V.; Jordan-Sciutto, K. L.; Hayley, S.; Mount, M. P.; O’Hare, M. J.; Callaghan, S.; Slack, R. S.; Przedborski, S.; Anisman, H.; Park, D. S., Cyclin-dependent kinase 5 is a mediator of dopaminergic neuron loss in a mouse model of Parkinson’s disease. Proc Natl Acad Sci U S A 2003, 100 (23), 13650–5. https://www.ncbi.nlm.nih.gov/pubmed/14595022.

68. Avraham, E.; Rott, R.; Liani, E.; Szargel, R.; Engelender, S., Phosphorylation of Parkin by the cyclin-dependent kinase 5 at the linker region modulates its ubiquitin-ligase activity and aggregation. J Biol Chem 2007, 282 (17), 12842–50. https://www.ncbi.nlm.nih.gov/pubmed/17327227.

69. Stenmark, H., Rab GTPases as coordinators of vesicle traffic. Nat Rev Mol Cell Biol 2009, 10 (8), 513–25. https://www.ncbi.nlm.nih.gov/pubmed/19603039.

70. Rankin, C. A.; Galeva, N. A.; Bae, K.; Ahmad, M. N.; Witte, T. M.; Richter, M. L., Isolated RING2 domain of parkin is sufficient for E2-dependent E3 ligase activity. Biochemistry 2014, 53 (1), 225–34. https://www.ncbi.nlm.nih.gov/pubmed/24328108.

71. Rath, S.; Sharma, R.; Gupta, R.; Ast, T.; Chan, C.; Durham, T. J.; Goodman, R. P.; Grabarek, Z.; Haas, M. E.; Hung, W. H. W.; Joshi, P. R.; Jourdain, A. A.; Kim, S. H.; Kotrys, A. V.; Lam, S. S.; McCoy, J. G.; Meisel, J. D.; Miranda, M.; Panda, A.; Patgiri, A.; Rogers, R.; Sadre, S.; Shah, H.; Skinner, O. S.; To, T. L.; Walker, M. A.; Wang, H.; Ward, P. S.; Wengrod, J.; Yuan, C. C.; Calvo, S. E.; Mootha, V. K., MitoCarta3.0: an updated mitochondrial proteome now with sub-organelle localization and pathway annotations. Nucleic Acids Res 2021, 49 (D1), D1541–D1547. https://www.ncbi.nlm.nih.gov/pubmed/33174596.

72. Scott, T. L.; Wicker, C. A.; Suganya, R.; Dhar, B.; Pittman, T.; Horbinski, C.; Izumi, T., Polyubiquitination of apurinic/apyrimidinic endonuclease 1 by Parkin. Mol Carcinog 2017, 56 (2), 325–336. https://www.ncbi.nlm.nih.gov/pubmed/27148961.

73. Sun, Y.; Vashisht, A. A.; Tchieu, J.; Wohlschlegel, J. A.; Dreier, L., Voltage-dependent anion channels (VDACs) recruit Parkin to defective mitochondria to promote mitochondrial autophagy. J Biol Chem 2012, 287 (48), 40652–60. https://www.ncbi.nlm.nih.gov/pubmed/23060438.

74. Bertolin, G.; Jacoupy, M.; Traver, S.; Ferrando-Miguel, R.; Saint Georges, T.; Grenier, K.; Ardila-Osorio, H.; Muriel, M. P.; Takahashi, H.; Lees, A. J.; Gautier, C.; Guedin, D.; Coge, F.; Fon, E. A.; Brice, A.; Corti, O., Parkin maintains mitochondrial levels of the protective Parkinson’s disease-related enzyme 17- beta hydroxysteroid dehydrogenase type 10. Cell Death Differ 2015, 22 (10), 1563–76. https://www.ncbi.nlm.nih.gov/pubmed/25591737.

75. Langemeyer, L.; Frohlich, F.; Ungermann, C., Rab GTPase Function in Endosome and Lysosome Biogenesis. Trends Cell Biol 2018, 28 (11), 957–970. https://www.ncbi.nlm.nih.gov/pubmed/30025982.

76. Borchers, A. C.; Langemeyer, L.; Ungermann, C., Who’s in control? Principles of Rab GTPase activation in endolysosomal membrane trafficking and beyond. J Cell Biol 2021, 220 (9). https://www.ncbi.nlm.nih.gov/pubmed/34383013.

77. Jimenez-Orgaz, A.; Kvainickas, A.; Nagele, H.; Denner, J.; Eimer, S.; Dengjel, J.; Steinberg, F., Control of RAB7 activity and localization through the retromer-TBC1D5 complex enables RAB7- dependent mitophagy. Embo J 2018, 37 (2), 235–254. https://www.ncbi.nlm.nih.gov/pubmed/29158324.

78. Yamano, K.; Wang, C.; Sarraf, S. A.; Munch, C.; Kikuchi, R.; Noda, N. N.; Hizukuri, Y.; Kanemaki, M. T.; Harper, W.; Tanaka, K.; Matsuda, N.; Youle, R. J., Endosomal Rab cycles regulate Parkin-mediated mitophagy. Elife 2018, 7. https://www.ncbi.nlm.nih.gov/pubmed/29360040.

79. Yamano, K.; Fogel, A. I.; Wang, C.; van der Bliek, A. M.; Youle, R. J., Mitochondrial Rab GAPs govern autophagosome biogenesis during mitophagy. Elife 2014, 3, e01612. https://www.ncbi.nlm.nih.gov/pubmed/24569479.

80. Tremel, S.; Ohashi, Y.; Morado, D. R.; Bertram, J.; Perisic, O.; Brandt, L. T. L.; von Wrisberg, M. K.; Chen, Z. A.; Maslen, S. L.; Kovtun, O.; Skehel, M.; Rappsilber, J.; Lang, K.; Munro, S.; Briggs, J. A. G.; Williams, R. L., Structural basis for VPS34 kinase activation by Rab1 and Rab5 on membranes. Nat Commun 2021, 12 (1), 1564. https://www.ncbi.nlm.nih.gov/pubmed/33692360.

81. Webster, C. P.; Smith, E. F.; Bauer, C. S.; Moller, A.; Hautbergue, G. M.; Ferraiuolo, L.; Myszczynska, M. A.; Higginbottom, A.; Walsh, M. J.; Whitworth, A. J.; Kaspar, B. K.; Meyer, K.; Shaw, P. J.; Grierson, A. J.; De Vos, K. J., The C9orf72 protein interacts with Rab1a and the ULK1 complex to regulate initiation of autophagy. Embo J 2016, 35 (15), 1656–76. https://www.ncbi.nlm.nih.gov/pubmed/27334615.

82. Martinez-Menarguez, J. A.; Martinez-Alonso, E.; Cara-Esteban, M.; Tomas, M., Focus on the Small GTPase Rab1: A Key Player in the Pathogenesis of Parkinson’s Disease. Int J Mol Sci 2021, 22 (21). https://www.ncbi.nlm.nih.gov/pubmed/34769517.

83. Wauters, F.; Cornelissen, T.; Imberechts, D.; Martin, S.; Koentjoro, B.; Sue, C.; Vangheluwe, P.; Vandenberghe, W., LRRK2 mutations impair depolarization-induced mitophagy through inhibition of mitochondrial accumulation of RAB10. Autophagy 2020, 16 (2), 203–222. https://www.ncbi.nlm.nih.gov/pubmed/30945962.

84. Jiang, P.; Nishimura, T.; Sakamaki, Y.; Itakura, E.; Hatta, T.; Natsume, T.; Mizushima, N., The HOPS complex mediates autophagosome-lysosome fusion through interaction with syntaxin 17. Mol Biol Cell 2014, 25 (8), 1327–37. https://www.ncbi.nlm.nih.gov/pubmed/24554770.

85. Hammerling, B. C.; Najor, R. H.; Cortez, M. Q.; Shires, S. E.; Leon, L. J.; Gonzalez, E. R.; Boassa, D.; Phan, S.; Thor, A.; Jimenez, R. E.; Li, H.; Kitsis, R. N.; Dorn, G. W., II; Sadoshima, J.; Ellisman, M. H.; Gustafsson, A. B., A Rab5 endosomal pathway mediates Parkin-dependent mitochondrial clearance. Nat Commun 2017, 8, 14050. https://www.ncbi.nlm.nih.gov/pubmed/28134239.

86. Kinchen, J. M.; Ravichandran, K. S., Identification of two evolutionarily conserved genes regulating processing of engulfed apoptotic cells. Nature 2010, 464 (7289), 778–82. https://www.ncbi.nlm.nih.gov/pubmed/20305638.

87. Poteryaev, D.; Datta, S.; Ackema, K.; Zerial, M.; Spang, A., Identification of the switch in early-to-late endosome transition. Cell 2010, 141 (3), 497–508. https://www.ncbi.nlm.nih.gov/pubmed/20434987.

88. Stroupe, C., This Is the End: Regulation of Rab7 Nucleotide Binding in Endolysosomal Trafficking and Autophagy. Front Cell Dev Biol 2018, 6, 129. https://www.ncbi.nlm.nih.gov/pubmed/30333976.

89. Jin, X.; Wang, K.; Wang, L.; Liu, W.; Zhang, C.; Qiu, Y.; Liu, W.; Zhang, H.; Zhang, D.; Yang, Z.; Wu, T.; Li, J., RAB7 activity is required for the regulation of mitophagy in oocyte meiosis and oocyte quality control during ovarian aging. Autophagy 2022, 18 (3), 643–660. https://www.ncbi.nlm.nih.gov/pubmed/34229552.

90. Song, P.; Trajkovic, K.; Tsunemi, T.; Krainc, D., Parkin Modulates Endosomal Organization and Function of the Endo-Lysosomal Pathway. J Neurosci 2016, 36 (8), 2425–37. https://www.ncbi.nlm.nih.gov/pubmed/26911690.

91. Peng, W.; Schroder, L. F.; Song, P.; Wong, Y. C.; Krainc, D., Parkin regulates amino acid homeostasis at mitochondria-lysosome (M/L) contact sites in Parkinson’s disease. Sci Adv 2023, 9 (29), eadh3347. https://www.ncbi.nlm.nih.gov/pubmed/37467322.

92. Sapmaz, A.; Berlin, I.; Bos, E.; Wijdeven, R. H.; Janssen, H.; Konietzny, R.; Akkermans, J. J.; Erson-Bensan, A. E.; Koning, R. I.; Kessler, B. M.; Neefjes, J.; Ovaa, H., USP32 regulates late endosomal transport and recycling through deubiquitylation of Rab7. Nat Commun 2019, 10 (1), 1454. https://www.ncbi.nlm.nih.gov/pubmed/30926795.

93. Jung, J.; Baek, J.; Tae, K.; Shin, D.; Han, S.; Yang, W.; Yu, W.; Jung, S. M.; Park, S. H.; Choi, C. Y.; Lee, S., Structural mechanism for regulation of Rab7 by site-specific monoubiquitination. Int J Biol Macromol 2022, 194, 347–357. https://www.ncbi.nlm.nih.gov/pubmed/34801583.

94. Shin, D.; Na, W.; Lee, J. H.; Kim, G.; Baek, J.; Park, S. H.; Choi, C. Y.; Lee, S., Site-specific monoubiquitination downregulates Rab5 by disrupting effector binding and guanine nucleotide conversion. Elife 2017, 6. https://www.ncbi.nlm.nih.gov/pubmed/28968219.

95. Zimprich, A.; Biskup, S.; Leitner, P.; Lichtner, P.; Farrer, M.; Lincoln, S.; Kachergus, J.; Hulihan, M.; Uitti, R. J.; Calne, D. B.; Stoessl, A. J.; Pfeiffer, R. F.; Patenge, N.; Carbajal, I. C.; Vieregge, P.; Asmus, F.; Muller-Myhsok, B.; Dickson, D. W.; Meitinger, T.; Strom, T. M.; Wszolek, Z. K.; Gasser, T., Mutations in LRRK2 cause autosomal-dominant parkinsonism with pleomorphic pathology. Neuron 2004, 44 (4), 601–7. https://www.ncbi.nlm.nih.gov/pubmed/15541309.

96. West, A. B.; Moore, D. J.; Biskup, S.; Bugayenko, A.; Smith, W. W.; Ross, C. A.; Dawson, V. L.; Dawson, T. M., Parkinson’s disease-associated mutations in leucine-rich repeat kinase 2 augment kinase activity. Proc Natl Acad Sci U S A 2005, 102 (46), 16842–7. https://www.ncbi.nlm.nih.gov/pubmed/16269541.

97. Steger, M.; Tonelli, F.; Ito, G.; Davies, P.; Trost, M.; Vetter, M.; Wachter, S.; Lorentzen, E.; Duddy, G.; Wilson, S.; Baptista, M. A.; Fiske, B. K.; Fell, M. J.; Morrow, J. A.; Reith, A. D.; Alessi, D. R.; Mann, M., Phosphoproteomics reveals that Parkinson’s disease kinase LRRK2 regulates a subset of Rab GTPases. Elife 2016, 5. https://www.ncbi.nlm.nih.gov/pubmed/26824392.

98. Vides, E. G.; Adhikari, A.; Chiang, C. Y.; Lis, P.; Purlyte, E.; Limouse, C.; Shumate, J. L.; Spinola- Lasso, E.; Dhekne, H. S.; Alessi, D. R.; Pfeffer, S. R., A feed-forward pathway drives LRRK2 kinase membrane recruitment and activation. Elife 2022, 11. https://www.ncbi.nlm.nih.gov/pubmed/36149401.

99. Eguchi, T.; Kuwahara, T.; Sakurai, M.; Komori, T.; Fujimoto, T.; Ito, G.; Yoshimura, S. I.; Harada, A.; Fukuda, M.; Koike, M.; Iwatsubo, T., LRRK2 and its substrate Rab GTPases are sequentially targeted onto stressed lysosomes and maintain their homeostasis. Proc Natl Acad Sci U S A 2018, 115 (39), E9115–E9124. https://www.ncbi.nlm.nih.gov/pubmed/30209220.

100. Bonet-Ponce, L.; Beilina, A.; Williamson, C. D.; Lindberg, E.; Kluss, J. H.; Saez-Atienzar, S.; Landeck, N.; Kumaran, R.; Mamais, A.; Bleck, C. K. E.; Li, Y.; Cookson, M. R., LRRK2 mediates tubulation and vesicle sorting from lysosomes. Sci Adv 2020, 6 (46). https://www.ncbi.nlm.nih.gov/pubmed/33177079.

101. Steger, M.; Diez, F.; Dhekne, H. S.; Lis, P.; Nirujogi, R. S.; Karayel, O.; Tonelli, F.; Martinez, T. N.; Lorentzen, E.; Pfeffer, S. R.; Alessi, D. R.; Mann, M., Systematic proteomic analysis of LRRK2-mediated Rab GTPase phosphorylation establishes a connection to ciliogenesis. Elife 2017, 6. https://www.ncbi.nlm.nih.gov/pubmed/29125462.

102. Dhekne, H. S.; Yanatori, I.; Gomez, R. C.; Tonelli, F.; Diez, F.; Schule, B.; Steger, M.; Alessi, D. R.; Pfeffer, S. R., A pathway for Parkinson’s Disease LRRK2 kinase to block primary cilia and Sonic hedgehog signaling in the brain. Elife 2018, 7. https://www.ncbi.nlm.nih.gov/pubmed/30398148.

103. Di Maio, R.; Hoffman, E. K.; Rocha, E. M.; Keeney, M. T.; Sanders, L. H.; De Miranda, B. R.; Zharikov, A.; Van Laar, A.; Stepan, A. F.; Lanz, T. A.; Kofler, J. K.; Burton, E. A.; Alessi, D. R.; Hastings, T. G.; Greenamyre, J. T., LRRK2 activation in idiopathic Parkinson’s disease. Sci Transl Med 2018, 10 (451). https://www.ncbi.nlm.nih.gov/pubmed/30045977.

104. Fujita, K.; Kedashiro, S.; Yagi, T.; Hisamoto, N.; Matsumoto, K.; Hanafusa, H., The ULK complex- LRRK1 axis regulates Parkin-mediated mitophagy via Rab7 Ser-72 phosphorylation. J Cell Sci 2022, 135 (23). https://www.ncbi.nlm.nih.gov/pubmed/36408770.

105. Heo, J. M.; Ordureau, A.; Swarup, S.; Paulo, J. A.; Shen, K.; Sabatini, D. M.; Harper, J. W., RAB7A phosphorylation by TBK1 promotes mitophagy via the PINK-PARKIN pathway. Sci Adv 2018, 4 (11), eaav0443. https://www.ncbi.nlm.nih.gov/pubmed/30627666.

106. Heo, J. M.; Ordureau, A.; Paulo, J. A.; Rinehart, J.; Harper, J. W., The PINK1-PARKIN Mitochondrial Ubiquitylation Pathway Drives a Program of OPTN/NDP52 Recruitment and TBK1 Activation to Promote Mitophagy. Mol Cell 2015, 60 (1), 7–20. https://www.ncbi.nlm.nih.gov/pubmed/26365381.

107. Ng, E. L.; Tang, B. L., Rab GTPases and their roles in brain neurons and glia. Brain Res Rev 2008, 58 (1), 236–46. https://www.ncbi.nlm.nih.gov/pubmed/18485483.

108. Shikanai, M.; Yuzaki, M.; Kawauchi, T., Rab family small GTPases-mediated regulation of intracellular logistics in neural development. Histol Histopathol 2018, 33 (8), 765–771. https://www.ncbi.nlm.nih.gov/pubmed/29266163.

109. Cortese, G. P.; Zhu, M.; Williams, D.; Heath, S.; Waites, C. L., Parkin Deficiency Reduces Hippocampal Glutamatergic Neurotransmission by Impairing AMPA Receptor Endocytosis. J Neurosci 2016, 36 (48), 12243–12258. https://www.ncbi.nlm.nih.gov/pubmed/27903732.

110. Helton, T. D.; Otsuka, T.; Lee, M. C.; Mu, Y.; Ehlers, M. D., Pruning and loss of excitatory synapses by the parkin ubiquitin ligase. Proc Natl Acad Sci U S A 2008, 105 (49), 19492–7. https://www.ncbi.nlm.nih.gov/pubmed/19033459.

111. Jiang, H.; Jiang, Q.; Feng, J., Parkin increases dopamine uptake by enhancing the cell surface expression of dopamine transporter. J Biol Chem 2004, 279 (52), 54380–6. https://www.ncbi.nlm.nih.gov/pubmed/15492001.

112. Zhu, M.; Cortese, G. P.; Waites, C. L., Parkinson’s disease-linked Parkin mutations impair glutamatergic signaling in hippocampal neurons. BMC Biol 2018, 16 (1), 100. https://www.ncbi.nlm.nih.gov/pubmed/30200940.

113. Kawabe, H.; Stegmuller, J., The role of E3 ubiquitin ligases in synapse function in the healthy and diseased brain. Mol Cell Neurosci 2021, 112, 103602. https://www.ncbi.nlm.nih.gov/pubmed/33581237.

114. Song, P.; Peng, W.; Sauve, V.; Fakih, R.; Xie, Z.; Ysselstein, D.; Krainc, T.; Wong, Y. C.; Mencacci, N. E.; Savas, J. N.; Surmeier, D. J.; Gehring, K.; Krainc, D., Parkinson’s disease-linked parkin mutation disrupts recycling of synaptic vesicles in human dopaminergic neurons. Neuron 2023, 111 (23), 3775–3788 e7. https://www.ncbi.nlm.nih.gov/pubmed/37716354.

115. Zhang, Y.; Gao, J.; Chung, K. K.; Huang, H.; Dawson, V. L.; Dawson, T. M., Parkin functions as an E2-dependent ubiquitin- protein ligase and promotes the degradation of the synaptic vesicle-associated protein, CDCrel-1. Proc Natl Acad Sci U S A 2000, *97* (24), 13354-9. https://www.ncbi.nlm.nih.gov/pubmed/11078524.

116. Trempe, J. F.; Chen, C. X.; Grenier, K.; Camacho, E. M.; Kozlov, G.; McPherson, P. S.; Gehring, K.; Fon, E. A., SH3 domains from a subset of BAR proteins define a Ubl-binding domain and implicate parkin in synaptic ubiquitination. Mol Cell 2009, 36 (6), 1034–47. https://www.ncbi.nlm.nih.gov/pubmed/20064468.

117. Bakker, J.; Spits, M.; Neefjes, J.; Berlin, I., The EGFR odyssey - from activation to destruction in space and time. J Cell Sci 2017, 130 (24), 4087–4096. https://www.ncbi.nlm.nih.gov/pubmed/29180516.

118. Mendoza, P.; Ortiz, R.; Diaz, J.; Quest, A. F.; Leyton, L.; Stupack, D.; Torres, V. A., Rab5 activation promotes focal adhesion disassembly, migration and invasiveness in tumor cells. J Cell Sci 2013, 126 (Pt 17), 3835–47. https://www.ncbi.nlm.nih.gov/pubmed/23813952.

119. Williams, K. C.; Coppolino, M. G., Phosphorylation of membrane type 1-matrix metalloproteinase (MT1-MMP) and its vesicle-associated membrane protein 7 (VAMP7)-dependent trafficking facilitate cell invasion and migration. J Biol Chem 2011, 286 (50), 43405–16. https://www.ncbi.nlm.nih.gov/pubmed/22002060.

120. Zhao, T.; Ding, X.; Yan, C.; Du, H., Endothelial Rab7 GTPase mediates tumor growth and metastasis in lysosomal acid lipase-deficient mice. J Biol Chem 2017, 292 (47), 19198–19208. https://www.ncbi.nlm.nih.gov/pubmed/28924047.

121. Ding, X.; Zhang, W.; Zhao, T.; Yan, C.; Du, H., Rab7 GTPase controls lipid metabolic signaling in myeloid-derived suppressor cells. Oncotarget 2017, 8 (18), 30123–30137. https://www.ncbi.nlm.nih.gov/pubmed/28415797.

122. Ioannou, M. S.; McPherson, P. S., Regulation of Cancer Cell Behavior by the Small GTPase Rab13. J Biol Chem 2016, 291 (19), 9929–37. https://www.ncbi.nlm.nih.gov/pubmed/27044746.

123. Ao, C.; Li, C.; Chen, J.; Tan, J.; Zeng, L., The role of Cdk5 in neurological disorders. Front Cell Neurosci 2022, 16, 951202. https://www.ncbi.nlm.nih.gov/pubmed/35966199.

124. Yan, J.; Zhang, P.; Tan, J.; Li, M.; Xu, X.; Shao, X.; Fang, F.; Zou, Z.; Zhou, Y.; Tian, B., Cdk5 phosphorylation-induced SIRT2 nuclear translocation promotes the death of dopaminergic neurons in Parkinson’s disease. NPJ Parkinsons Dis 2022, 8 (1), 46. https://www.ncbi.nlm.nih.gov/pubmed/35443760.

125. Shukla, A. K.; Spurrier, J.; Kuzina, I.; Giniger, E., Hyperactive Innate Immunity Causes Degeneration of Dopamine Neurons upon Altering Activity of Cdk5. Cell Rep 2019, 26 (1), 131–144 e4. https://www.ncbi.nlm.nih.gov/pubmed/30605670.

126. Conway, P.; Tyka, M. D.; DiMaio, F.; Konerding, D. E.; Baker, D., Relaxation of backbone bond geometry improves protein energy landscape modeling. Protein Sci 2014, 23 (1), 47–55. <GO to ISI>://WOS:000328566600004.

127. Barbas, C. F., 3rd; Burton, D. R.; Scott, J. K.; Silverman, G. J., Phage Display, A laboratory Manual. Cold Spring Harbor Laboratory Press: New York, 2000.

128. Kay, B. K.; Winter, J.; McCafferty, J., Phage Display of Peptides and Proteins. Academic Press, Inc.: Boston, 1996.

129. Perez-Riverol, Y.; Csordas, A.; Bai, J.; Bernal-Llinares, M.; Hewapathirana, S.; Kundu, D. J.; Inuganti, A.; Griss, J.; Mayer, G.; Eisenacher, M.; Perez, E.; Uszkoreit, J.; Pfeuffer, J.; Sachsenberg, T.; Yilmaz, S.; Tiwary, S.; Cox, J.; Audain, E.; Walzer, M.; Jarnuczak, A. F.; Ternent, T.; Brazma, A.; Vizcaino, J. A., The PRIDE database and related tools and resources in 2019: improving support for quantification data. Nucleic Acids Res 2019, 47 (D1), D442–D450. https://www.ncbi.nlm.nih.gov/pubmed/30395289.

